# Neuronal Switching Between Single- and Dual-Network Activity via Modulation of Intrinsic Membrane Properties

**DOI:** 10.1101/2021.02.05.429848

**Authors:** Savanna-Rae H. Fahoum, Dawn M. Blitz

## Abstract

Oscillatory networks underlie rhythmic behaviors (e.g. walking, chewing), and complex behaviors (e.g. memory formation, decision making). Flexibility of oscillatory networks includes neurons switching between single- and dual-network participation, even generating oscillations at two distinct frequencies. Modulation of synaptic strength can underlie this neuronal switching. Here we ask whether switching into dual-frequency oscillations can also result from modulation of intrinsic neuronal properties. The isolated stomatogastric nervous system of male Cancer borealis crabs contains two well-characterized rhythmic feeding-related networks (pyloric, ∼1 Hz; gastric mill, ∼0.1 Hz). The identified modulatory projection neuron MCN5 causes the pyloric-only LPG neuron to switch to dual pyloric/gastric mill bursting. Bath applying the MCN5 neuropeptide transmitter Gly^1^-SIFamide only partly mimics the LPG switch to dual activity, due to continued LP neuron inhibition of LPG. Here, we find that MCN5 uses a co-transmitter, glutamate, to inhibit LP, unlike Gly^1^-SIFamide excitation of LP. Thus, we modeled the MCN5-elicited LPG switching with Gly^1^-SIFamide application and LP photoinactivation. Using hyperpolarization of pyloric pacemaker neurons and gastric mill network neurons, we found that LPG pyloric-timed oscillations require rhythmic electrical synaptic input. However, LPG gastric mill-timed oscillations do not require any pyloric/gastric mill synaptic input and are voltage dependent. Thus, we identify modulation of intrinsic properties as an additional mechanism for switching a neuron into dual-frequency activity. Instead of synaptic modulation switching a neuron into a second network as a passive follower, modulation of intrinsic properties could enable a switching neuron to become an active contributor to rhythm generation in the second network.

**Significance Statement:** Neuromodulation of oscillatory networks can enable network neurons to switch from sing<bacle- to dual-network participation, even when two networks oscillate at distinct frequencies. We used small, well-characterized networks to determine whether modulation of synaptic strength, an identified mechanism for switching, is necessary for dual-network recruitment. We demonstrate that rhythmic electrical synaptic input is required for continued linkage with a “home” network, but that modulation of intrinsic properties is sufficient to switch a neuron into dual-frequency oscillations, linking it to a second network. Neuromodulator-induced switches in neuronal participation between networks occurs in motor, cognitive, and sensory networks. Our study highlights the importance of considering intrinsic properties as a pivotal target for enabling parallel participation of a neuron in two oscillatory networks.

## Introduction

Oscillatory networks are important for rhythmic motor behaviors such as locomotion and complex behaviors such as memory processing (McCormick and Bal, 1997; Buzsáki, 2002; Wang, 2010; Bucher et al., 2015). Individual neurons can participate in more than one rhythmic network, and even oscillate at multiple frequencies simultaneously. This includes neurons bursting in time with breathing and vocalizing, or other orofacial networks, with multiple feeding networks, and with multiple hippocampal or cortical rhythms (Steriade et al., 1993; Dickinson, 1995; Roopun et al., 2008; Bartlett and Leiter, 2012). However, multiplexing of neuronal activity across networks is not a static feature.

Network flexibility extends to neurons switching participation between networks (Dickinson, 1995; Bouret and Sara, 2005; Koch et al., 2011). This “neuronal switching” includes neurons being recruited into or removed from a network; neurons from multiple networks joining together into a single, novel network; or neurons switching between single- and dual-network participation (Dickinson et al., 1990; Hooper and Moulins, 1990; Meyrand et al., 1994; Weimann and Marder, 1994; Tryba et al., 2008; Koch et al., 2011). Small invertebrate motor systems enable identification of modulatory projection neurons as endogenous sources of neuromodulators that elicit neuronal switching (Meyrand et al., 1994; Faumont et al., 2005). Based on invertebrate and systems-level vertebrate studies, it is proposed that modulatory neurons in vertebrate nervous systems similarly elicit neuronal switching (Bouret and Sara, 2005). For example, exogenous modulators elicit switching in respiratory networks (Tryba et al., 2008), noradrenergic signaling regulates coupling of brain regions during stress responses in human fMRI measurements, and locus coeruleus activation elicits cognitive shifts and large-scale connectivity changes in rodents and monkeys (Bouret and Sara, 2005; Hermans et al., 2011; Zerbi et al., 2019). However, it is difficult to selectively manipulate modulatory neurons and identify the cellular mechanisms of neuromodulator-elicited neuronal switching in large diffuse networks.

In general, neuromodulators alter the intrinsic membrane properties of neurons and/or the synapses between them (Harris-Warrick, 2011; Marder et al., 2014; Nadim and Bucher, 2014). In invertebrate networks, modulation of synaptic strength is necessary for recruiting switching neurons between networks (Hooper and Moulins 1990; Meyrand et al. 1994). Although work in larger vertebrate networks and computational studies suggest that modulation of intrinsic membrane properties also contributes to switching (Tryba et al., 2008; Drion et al., 2019), complications such as electrical coupling have prevented confirming this in a biological context. Further, it remains unknown whether intrinsic properties can be sufficient to enable neuronal switching into a second oscillation frequency.

To study neuronal switching, we utilized the isolated stomatogastric nervous system (STNS) of the crab *C. borealis*, which contains identified modulatory inputs that alter well-defined networks (Nusbaum and Beenhakker, 2002; Stein, 2009; Marder, 2012; Daur et al., 2016). The STNS includes two networks totaling 26-30 neurons that generate the pyloric (food filtering, ∼1 Hz) and gastric mill (food chewing, ∼0.1 Hz) rhythms (Nusbaum and Beenhakker, 2002; Daur et al., 2016) (Fig. 1). Most of these neurons can participate in both feeding networks, depending on their modulatory state (Bucher et al. 2006; Dickinson et al. 1990; Hooper and Moulins 1990; Meyrand et al. 1994; Weimann et al. 1991). Furthermore, neuromodulatory input-elicited neuronal switching is well-established in the STNS (Hooper and Moulins 1990; Meyrand et al. 1994).

**Figure 1.**
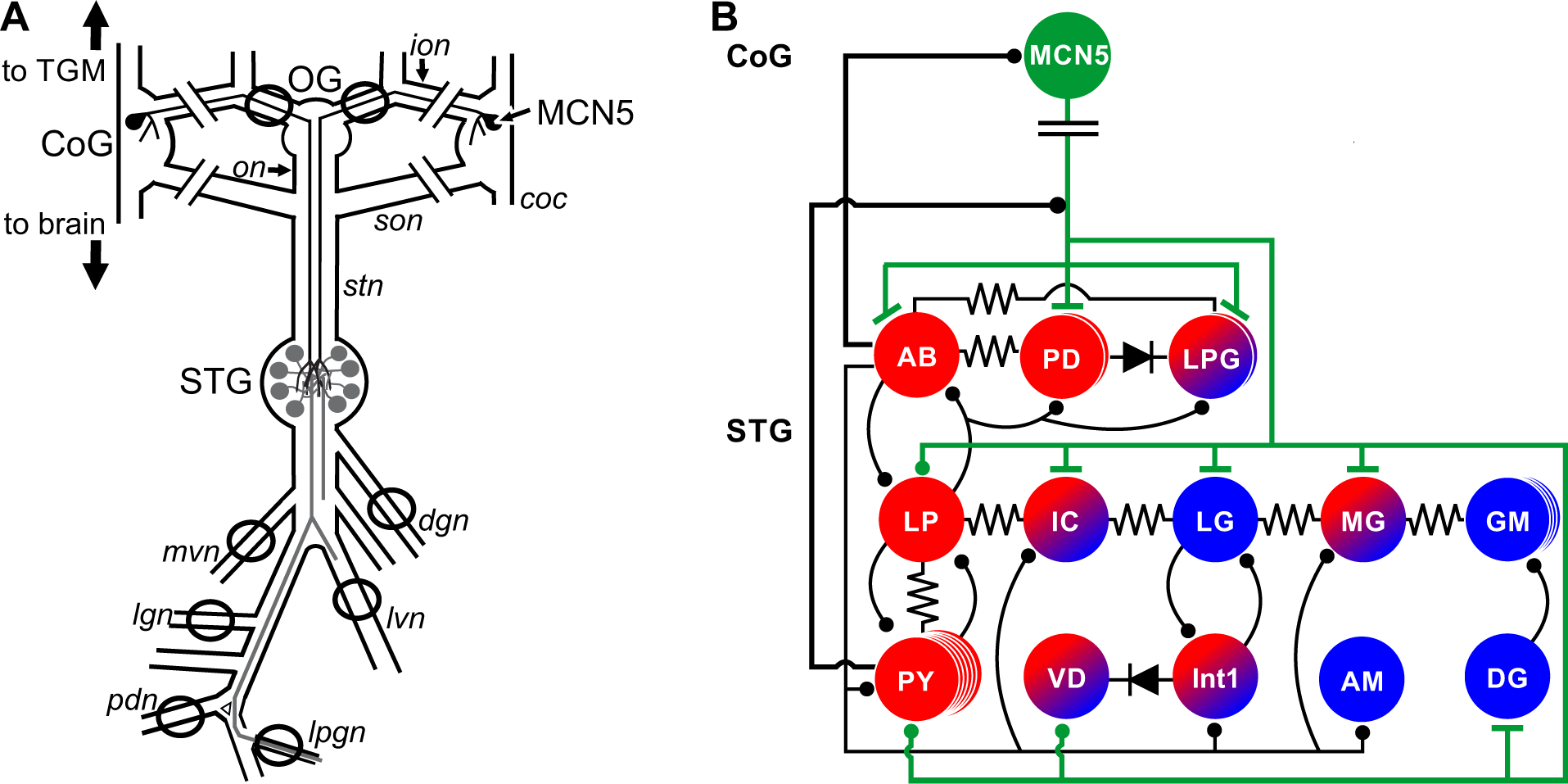
Schematic of the isolated stomatogastric nervous system and connectivity of pyloric and gastric mill networks including input from a modulatory projection neuron, MCN5. *A*, The isolated stomatogastric nervous system is comprised of the paired commissural ganglia (CoG), oesophageal ganglion (OG), and stomatogastric ganglion (STG), as well as their connecting and peripheral nerves. A single modulatory commissural neurons 5 (MCN5) projects from each CoG through the *ion* and *stn* to the STG (Norris et al., 1996). The line breaks on the *sons* and *ions* indicate where these nerves were cut to isolate the STG networks (see Methods). *B*, The related pyloric and gastric mill networks have extensive electrical (resistor and diode symbols) and chemical inhibitory (ball and stick) connections between network neurons. Some neurons are exclusively active with the pyloric network (red), some exclusively with the gastric mill network (blue) and some can be active with both networks (red/blue) under different modulatory conditions. MCN5 inhibits the PY and LP neurons (ball and stick), and through modulatory actions, excites the pyloric pacemaker ensemble (AB, PD, LPG; t-bar symbols), and activates the gastric mill neurons IC, LG, and DG (Norris et al., 1996; Blitz et al., 2019). The line break in the MCN5 axon indicate additional distance between the CoGs and STG. MCN5 connectivity is shown in green; thick black lines from AB and PY are feedback to MCN5; thin black lines are synaptic connections within the STG. Abbreviations: Ganglia: CoG, commissural ganglion; OG: oesophageal ganglion; STG stomatogastric ganglion; Nerves: *dgn*, dorsal gastric nerve; *lgn*, lateral gastric nerve; *lpgn*, lateral posterior gastric nerve; *lvn*, lateral ventricular nerve; *mvn*, medial ventricular nerve; *pdn*, pyloric dilator nerve; Neurons: AB, anterior burster; AM, anterior medial; DG, dorsal gastric; GM, gastric mill; IC, inferior cardiac; Int1, interneuron 1; LG, lateral gastric; LP, lateral pyloric; LPG, lateral posterior gastric; MG, medial gastric; PD, pyloric dilator; PY, pyloric; VD, ventricular dilator.

In *C. borealis*, modulatory commissural neuron 5 (MCN5) elicits a novel rhythm in which the pyloric-only, lateral posterior gastric (LPG) neuron switches participation to dual pyloric/gastric mill bursting (Norris et al., 1996; Blitz et al., 2019). During MCN5 activation, slower long-duration, gastric mill-timed bursts are superimposed on the faster pyloric-timed LPG bursting, despite strong electrical coupling to the pyloric pacemaker neurons (Blitz et al., 2019). Here, we examined whether electrical and/or chemical synaptic input is necessary for LPG to switch between single- and dual-network activity. We found that: 1) both bath-applied and neuronally-released neuropeptide elicit neuronal switching via modulation of intrinsic properties; 2) electrical synaptic input works in concert with intrinsic properties to generate dual-frequency activity.

## Materials and Methods

### Animals

Adult male *C. borealis* crabs were obtained from The Fresh Lobster Company (Gloucester, MA) and maintained in artificial seawater at 10-12 °C until used for experiments. Crabs were dissected to isolate the stomatogastric nervous system (STNS) (Gutierrez and Grashow, 2009; Blitz et al., 2019). Briefly, crabs were cold anesthetized by packing in ice for 35-50 min prior to dissection. The foregut was removed from the animal during gross dissection, bisected, and pinned flat in a Sylgard 170-lined dish (Fisher Scientific, Hampton, NH). During fine dissection, the STNS was carefully removed and pinned in a Sylgard 184-lined petri dish (Fisher Scientific). For all parts of the dissection, the preparation was kept in chilled (4 °C) *C. borealis* physiological saline.

### Solutions

*C. borealis* physiological saline was composed of (in mM): 440 NaCl, 26 MgCl2, 13 CaCl2, 11 KCl, 10 Trizma Base, 5 Maleic Acid, pH: 7.4 – 7.6. Squid internal electrode solution contained (in mM): 10 MgCl2, 400 potassium D-Gluconic Acid, 10 HEPES, 15 NaSO4, 20 NaCl, pH = 7.45. (Hooper et al., 2015). Gly^1^-SIFamide (GYRKPPFNG-SIFamide, custom peptide synthesis: Genscript, Piscataway, NJ) (Huybrechts et al., 2003; Yasuda et al., 2004; Dickinson et al., 2008; Blitz et al., 2019) was dissolved in optima water (Fisher Scientific, Waltham, MA) at 10^-2^ M and aliquots stored at -20 °C until needed. Gly^1^-SIFamide aliquots were diluted in physiological saline to a final concentration of 5 x 10^-6^ M. Picrotoxin (PTX) powder (Sigma Aldrich, St. Louis, MO) was added directly to physiological saline at a final concentration of 10^-5^ M and vigorously stirred for at least 45 min prior to use. For some experiments, Gly^1^-SIFamide aliquots were added directly to PTX in physiological saline (10^-5^ M) at a final concentration of 5 x 10^-6^ M.

### Electrophysiology

All preparations were continuously superfused with chilled *C. borealis* physiological saline (8-10 °C), or chilled saline containing Gly^1^-SIFamide and/or PTX as indicated. All solution changes were performed using a switching manifold for uninterrupted superfusion of the preparation. Using a Model 1700 A-M Systems Amplifier (A-M Systems, Carlsborg, WA), extracellular activity in nerves were recorded using custom-made stainless steel pin electrodes, with one wire placed in Vaseline wells that were built around each nerve, and one wire as reference outside the well. Stomatogastric ganglion (STG) somata were exposed by removing the thin layer of tissue across the ganglion and observed with light transmitted through a dark-field condenser (MBL-12010 Nikon Instruments, Inc, Melville, NY). Intracellular recordings of STG somata were obtained via sharp-tip glass microelectrodes (18-30 MΩ) filled with 0.6 M potassium acetate and 20 mM potassium chloride. For experiments involving intracellular LPG or MCN5 recordings, microelectrodes were filled with squid internal solution (20-40 MΩ) (see *Solutions*) which better maintains neuron properties over time (Hooper et al., 2015). All intracellular recordings were collected using AxoClamp 900A amplifiers in current clamp mode (Molecular Devices, San Jose, CA). All experiments, excluding those with intracellular MCN5 recordings, were performed in the isolated stomatogastric nervous system following transection of both inferior and superior oesophageal nerves (ion and son, respectively). For intracellular MCN5 recording experiments, the ipsilateral *ion* was not transected to retain the MCN5 projection to the STG. STG neurons were identified based on their nerve projection patterns and interactions with other STG neurons. All electrophysiological recordings were collected using acquisition hardware (Micro1401) and software (Spike2; ∼5 kHz sampling rate, Cambridge Electronic Design, Cambridge, UK) and laboratory computer (Dell, Round Rock, TX).

The pyloric neuron lateral pyloric (LP) and gastric mill neurons inferior cardiac (IC), dorsal gastric (DG), lateral gastric (LG), and medial gastric (MG) were hyperpolarized in order to eliminate spike-mediated and graded transmitter release. The current injection amplitude (-2 to -4 nA; 200 s) was sufficient to decrease their voltage oscillation amplitude indicating that the membrane potential was approaching reversal potential for inhibitory input and therefore below transmitter release threshold. The two pyloric dilator (PD) neurons were each hyperpolarized (-4 to -6 nA) to eliminate rhythmic oscillations of the pyloric pacemaker ensemble. Due to electrical coupling between anterior burster (AB) and the two PD neurons, hyperpolarization of the PD neurons hyperpolarizes AB, the main pacemaker in control conditions, and thus eliminates the pyloric rhythm (Bartos et al., 1999; Blitz et al., 2008). In PTX, to eliminate LP inhibition of the pacemakers and thus isolate the pacemaker influence on LPG, when both PDs were hyperpolarized, the most hyperpolarized LPG membrane potential did not change (PD On to Off: -51.5 ± 0.71 to -51.8 ± 0.91 mV, t(5) = 0.336, p = 0.751, n = 6, paired t-test), presumably due to the rectifying nature of this electrical coupling (Shruti et al., 2014). Each of the two PD neurons was hyperpolarized for 200 s, except when the hyperpolarization was maintained for the duration of PTX:Gly^1^-SIFamide applications to allow sufficient time to test the voltage-dependence of LPG oscillations. To determine whether LPG bursting was voltage-dependent, we injected hyperpolarizing and depolarizing current into LPG (± 0.5, 1, 1.5, 2 nA) until at least five full burst cycles occurred, or 200 s passed without a full burst cycle taking place. These experiments were conducted during Gly^1^-SIFamide in PTX with both PD neurons hyperpolarized to eliminate rhythmic pyloric electrical coupling input (PTX:PDhype:Gly^1^-SIFamide application).

### MCN5 identification and stimulation

MCN5 was stimulated via intracellular current injection, or extracellular *ion* stimulation. The MCN5 soma was identified by correlating its spike activity with spikes in the *ion*, and by its effects on STG neurons (Norris et al., 1996; Blitz et al., 2019). There are only two neurons that project from the CoG to the STG via the *ion*, MCN5 and modulatory commissural neuron 1 (MCN1) (Coleman and Nusbaum, 1994; Bartos and Nusbaum, 1997). MCN1 and MCN5 have distinct effects, including opposite actions on several STG neurons (Coleman and Nusbaum, 1994; Norris et al., 1996; Bartos and Nusbaum, 1997; Blitz et al., 2019). For extracellular MCN5 stimulation, the *ion* was stimulated at 30 Hz (3-6 V; Grass S88 stimulator and Grass SIU5 stimulus isolation unit; Grass Instruments) when MCN5 activation threshold was lower than that of MCN1, or following photoinactivation of the MCN1 axon in the stomatogastric nerve (stn) near the entrance to the STG (MCN1STG; see below) to eliminate MCN1 effects on STG neurons (Blitz et al., 2019).

### Photoinactivation

Photoinactivation is used to selectively eliminate neuron activity from a network without influencing electrically coupled neurons (Miller and Selverston, 1979; Marder and Eisen, 1984). The LP neuron or MCN1STG was impaled with a sharp microelectrode (25-40 MΩ) that was tip-filled with Alexa Fluor 568 hydrazide (10 mM in 200 mM KCl; Fisher Scientific) and backfilled with 0.6 M potassium acetate plus 20 mM potassium chloride. MCN1STG was identified based on the presence of 1:1 action potentials in MCN1STG during low-frequency (3 Hz) *ion* stimulation at MCN1 activation threshold, inhibitory post-synaptic potentials from the LG neuron, and corresponding electrical excitatory post-synaptic potentials (eEPSPs) in LG (Coleman and Nusbaum, 1994; Coleman et al., 1995). Hyperpolarizing current (-5 nA) was injected into LP for ∼30 minutes to fill the soma and neurites with the negatively charged Alexa 568 dye or into a small region of MCN1STG (-5 nA, ∼5 min). The STG or stn near its entrance to the STG was then illuminated for 3-7 min using a Texas red filter set (560 ± 40 nm wavelength; Leica, Wetzlar, Germany). Complete photoinactivation of LP was confirmed when the membrane potential reached 0 mV, and LP action potentials were absent from a lateral ventricular nerve (lvn) recording (Miller and Selverston, 1979). Photoinactivation of MCN1STG was confirmed when the membrane potential reached 0 mV and there were no longer MCN1-elicited eEPSPs in LG during *ion* stimulations.

### Data Analysis

All analyses were conducted after MCN5 or Gly^1^-SIFamide actions had reached a steady state. LP activity, including number of spikes per pyloric-timed burst and firing frequency ([number of spikes per burst – 1]/burst duration) was quantified across a 130-300 s window during sustained MCN5 stimulation or Gly^1^-SIFamide application. To test whether MCN5 uses glutamate to inhibit LP, we measured LP number of spikes per burst, firing frequency, and the most depolarized peak membrane potential, excluding action potentials, during rhythmic MCN5 stimulation (5 s duration, 30 Hz intraburst frequency, 0.1 Hz interburst frequency). LP parameters were measured across 5 s blocks of alternating MCN5 stimulation off and on. Data from 10 blocks per condition (10 off, 10 on), per experiment were averaged during saline and PTX application (10^-5^ M). In PTX, LP bursts were defined based on inhibition from PD neurons, as these neurons use cholinergic transmission to inhibit the LP neuron which is not blocked by PTX (Marder and Eisen, 1984).

### LPG Burst Identification

The LPG interspike interval (ISI) distribution was used to identify LPG pyloric- and gastric mill-timed bursts. First, a histogram of LPG ISIs across an entire MCN5 stimulation or Gly^1^-SIFamide application was generated and the two largest peaks were identified. The first of these peaks (within ∼0 – 0.5 s) included *intra*burst intervals (interval between spikes during a burst), and the second peak (∼0.5 – 2 s) included *inter*burst intervals (interval between spikes between bursts). We calculated the mean ISI between these two peaks and used this value as a cutoff, such that ISIs above the cutoff indicated the end of one and beginning of another burst. A cutoff ISI was determined in this manner for each preparation and was used to detect both pyloric and gastric mill-timed bursts for each experimental manipulation. The ISI cutoff value was initially calculated in Excel and then using a custom-written MATLAB (Mathworks, Natick, MA) function. To select only gastric mill-timed LPG bursts from all LPG bursts identified with the ISI cutoff, we used a custom-written Spike 2 script, that identifies LPG bursts with a duration greater than one pyloric cycle period (from PD neuron burst onset to the subsequent PD neuron burst onset; ∼1 s). Throughout the study, the number of these LPG gastric mill-timed bursts was quantified, as well as the LPG gastric mill-timed cycle period (duration between the first action potentials of two consecutive LPG gastric mill-timed bursts), burst duration (duration from the first to the last action potential in an LPG gastric mill-timed burst), and duty cycle (burst duration/cycle period) during MCN5 stimulation and Gly^1^-SIFamide application.

### Spectral analysis

We used spectral analysis to quantify the degree to which LPG participates in the pyloric and gastric mill rhythms (Bucher et al., 2006; Rehm et al., 2008). All spectral analysis was conducted on LPG activity recorded extracellularly from the lateral pyloric gastric nerve (*lpgn*) which includes activity of the two LPG neurons, which were always co-active. To calculate the LPG power spectrum, we measured LPG instantaneous spike frequency (IF = 1/ISI) across each analysis window and plotted these values on a histogram to distinguish between gastric mill (0.05 – 0.19 Hz) and pyloric (0.2 – 4.0 Hz) burst frequencies. The regions for gastric mill and pyloric activity were based on Bucher et al (2006), and the observed gastric mill-timed cycle periods of LPG bursting in preliminary Gly^1^-SIFamide application experiments. The count of LPG gastric mill- and pyloric-timed IFs within these two regions were summed and then normalized to the total IF count between 0.05 – 4.00 Hz for each manipulation in each preparation.

Normalized values were converted to a percentage to compare across preparations. Note that pyloric and gastric mill percentages add to 100%, therefore the amount of increase in one is the amount of decrease in the other and statistical results for pyloric and gastric mill activity yield the same values. Thus, only LPG gastric mill percentages were analyzed statistically and reported in the text.

To compare LPG bursts when isolated from both networks (10^-5^ M PTX:PDhype) to pyloric- and gastric mill-timed bursts in network-intact conditions, we quantified LPG burst parameters across five LPG gastric mill (slow) burst cycles, including all the pyloric-timed bursts within those five slow burst cycles in control and measured five cycles of isolated LPG bursts. In two preparations in which MCN5 was stimulated we were unable to analyze extracellular LPG activity with networks intact and thus analyzed LPG activity from an intracellular recording. To assess any voltage-dependence of LPG bursting during PTX:PDhype:Gly^1^-SIFamde application, we measured LPG cycle period and duty cycle across five burst cycles. For this data set, we analyzed the activity of the single intracellularly recorded LPG. We used ISI cutoff values (see above) to detect LPG bursts, however the ISI value was adjusted when spikes that were visually part of an LPG burst were not included with the ISI value (i.e., spikes that occurred at the end of a burst before the membrane potential returned to the trough potential). Additionally, detected bursts that were shorter than an average pyloric cycle (1 s) were excluded from analysis. LPG activity was categorized as tonic when the ISI cutoff value detected no interburst intervals and therefore no bursts, for the duration of the current injection, and as off when there were no action potential bursts.

### Software and Statistical Analysis

Raw data were analyzed using scripts/functions written in Spike 2 or MATLAB. ISI histograms were made in Excel (Microsoft, Redmond, WA) or MATLAB. Statistical analysis was performed with SigmaPlot (Systat, San Jose, CA). Final figures were made using CorelDraw (Corel, Ottawa, ON, Canada). Graphs were plotted in SigmaPlot or MATLAB and imported into CorelDraw. Data was first analyzed for normality to determine whether a parametric or non-parametric test was to be used on each data set. Paired t-test, Student’s t-test, Friedman One-way ANOVA on Ranks, Friedman One-way repeated measures (RM) ANOVA on Ranks, One-way RM ANOVA and post-hoc tests for multiple comparisons were used as indicted. Threshold for significance was p < 0.05. All data are presented as mean ± SE.

### Code Accessibility

Spike2 scripts and Matlab functions used for analysis are available upon request. Please contact the corresponding author.

## Results

The STG within the STNS receives modulatory input from the bilateral commissural ganglia (CoGs) and the oesophageal ganglion (OG) (Fig. 1A). The modulatory projection neuron, MCN5 soma is located in the CoG (one copy in each CoG), with axonal projections to the STG via the *ion* and *stn* (Fig. 1A). The pyloric (red) and gastric mill (blue) network neurons within the STG are connected by inhibitory chemical, and electrical synapses (Fig. 1B). The pyloric network is constitutively active in vivo and in vitro (Marder and Bucher, 2007; Stein, 2009), while the gastric mill network requires activation of modulatory projection or sensory neurons (Beenhakker et al., 2004; Blitz et al., 2004, 2008; Stein, 2009; Hedrich et al., 2011). When the gastric mill network is activated, some pyloric neuron activity becomes time-locked to both rhythms (red/blue) (Fig. 1B). MCN5 has classical and modulatory actions on most STG neurons and receives both long-distance and local presynaptic feedback from network neurons AB and pyloric (PY) neurons, respectively (Fig. 1B) (Norris et al., 1996; Blitz et al., 2019).

Tonic MCN5 stimulation (30 Hz) alters the activity of both STG networks (Norris et al., 1996; Blitz et al., 2019) (Fig. 2A). In particular, the pyloric rhythm increases in frequency, evident in the increased frequency of PD neuron (*pdn*) bursts (Fig 2A) (Norris et al., 1996; Blitz et al., 2019). Furthermore, a gastric mill rhythm is activated, indicated by the activation of rhythmic LG (*lgn*), DG (*dgn*), and IC neuron activity (Fig 2A) (Blitz et al., 2019). In addition, MCN5 decreases activity in the LP neuron (Fig 2A), and switches LPG neuron (*lpgn*) participation from pyloric only (Fig 2A, left) to dual pyloric (black arrows) and gastric mill (white arrows) network bursting (Fig. 2A, right) (Norris et al., 1996; Blitz et al., 2019). We refer to this LPG switch to dual pyloric (fast)/gastric mill (slow) bursting as dual-network activity.

**Figure 2.**
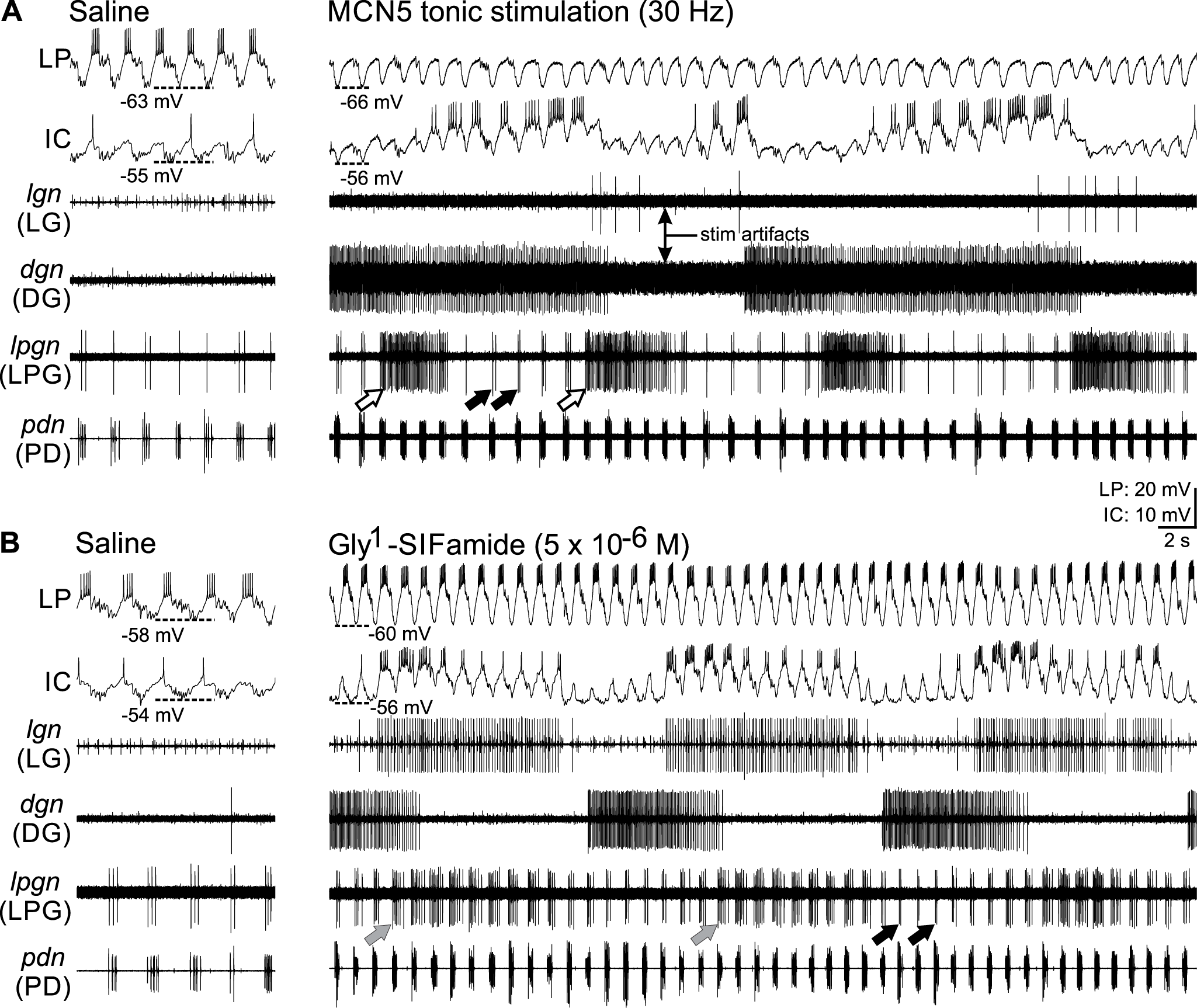
The MCN5 transmitter Gly^1^-SIFamide does not entirely mimic the effects of MCN5 stimulation. *A*, (*Left*) In the control condition prior to MCN5 stimulation, the pyloric rhythm was active evident in rhythmic bursting in the pyloric pacemaker neurons PD (*pdn*) and LPG (*lpgn*), and the LP neuron. The IC neuron was weakly active with 0-1 spikes per pyloric cycle. Gastric mill neurons DG and LG were silent. (*Right*) During tonic MCN5 stimulation (30 Hz), the pyloric rhythm was enhanced, evident by stronger, faster PD neuron bursting, and a gastric mill rhythm was activated, evident in the LG and DG neuron bursting (*lgn, dgn*, respectively). Additionally, LP neuron was inhibited, IC was active in time with both rhythms, and the LPG neuron switched to dual pyloric (*black arrows*) and gastric mill (*white arrows*) timed bursting. B, (*Left*) Again in control saline conditions the pyloric rhythm was weakly active (PD, LPG, LP, IC) and the gastric mill rhythm was silent (LG, DG). (Right). Bath application of Gly^1^-SIFamide (5 x 10^-6^ M) strengthened LP neuron activity, activated gastric mill network neurons LG (*lgn*), and DG (*dgn*), increased IC activity which was linked to both the pyloric and gastric mill rhythms, and increased the pyloric frequency (*pdn*). LPG (*lpgn*) did not switch to dual bursting, however, LPG exhibited periodic extensions of burst duration (grey arrows) for several pyloric cycles that coordinated with the gastric mill rhythm. (*A*) and (*B*) are from the same preparation.

MCN5 releases a peptide co-transmitter, Gly^1^-SIFamide (Blitz et al., 2019). Bath application of Gly^1^-SIFamide (5 x 10^-6^ M) partially mimics neuronal release from MCN5 (Blitz et al., 2019) (Fig. 2A-B). Similar to tonic MCN5 stimulation, Gly^1^-SIFamide application elicits an increase in pyloric frequency and activates a gastric mill rhythm (Blitz et al., 2019) (Fig. 2B, pyloric: pdn, gastric mill: *dgn, lgn*, IC). However, a key distinction between MCN5 and Gly^1^-SIFamide is in the LPG neuron response (Fig. 2B, *lpgn*). In control saline conditions, LPG is active only in pyloric time and coincident with PD neuron activity due to electrical coupling among the pyloric pacemaker neurons, AB, PD, and LPG (Marder and Eisen, 1984; Shruti et al., 2014) (Fig. 2B, left traces). During Gly^1^-SIFamide (5 x 10^-6^ M) application, LPG continues to generate pyloric-timed bursts (*lpgn, black arrows*), as well as periodic, longer-duration bursts in which LPG activity extends beyond a single PD burst (*lpgn*, grey arrows), but does not persist for multiple pyloric cycles (Blitz et al., 2019) (Fig. 2B). This is in contrast to the gastric mill timed LPG bursts during MCN5 stimulation, which persist for multiple pyloric cycles (Blitz et al., 2019) (Fig. 2A, white arrows).

In order to extend previous results (Blitz et al., 2019) and quantitatively compare LPG activity during MCN5 stimulation and Gly^1^-SIFamide application, we developed criteria to identify gastric-mill timed bursts. Specifically, LPG bursts were identified based on a minimum interburst interspike interval, and a minimum burst duration that was longer than one full pyloric cycle (see Methods). Using these criteria, four LPG slow bursts occurred during the example MCN5 stimulation (tonic 30 Hz; Fig. 2A, right), whereas zero LPG bursts met the criteria for a slow burst in an example Gly^1^-SIFamide application (Fig. 2B, right). Across preparations, LPG generated more slow bursts during MCN5 stimulation versus Gly^1^-SIFamide application within 200 s analysis windows (MCN5: 13 ± 2.1, n = 11; Gly^1^-SIFamide: 4.7 ± 1.2, n = 27; p < 0.001, Mann-Whitney Rank Sum Test). Thus, our quantification aligns with the previous qualitative assessment (Blitz et al., 2019), that Gly^1^-SIFamide does not entirely mimic the MCN5-elicited LPG switch from single-to dual-network activity.

Gly^1^-SIFamide and LP photoinactivation to model MCN5 stimulation To determine why Gly^1^-SIFamide did not entirely mimic the MCN5 actions on LPG, we considered the well-described circuitry of the pyloric and gastric mill networks, as well as other Gly^1^-SIFamide effects. LPG receives two inputs from STG neurons: 1) feedback inhibition from the LP neuron, and 2) electrical synaptic input from the pyloric pacemaker ensemble (Marder and Eisen, 1984; Shruti et al., 2014) (Fig. 1B). Similar to MCN5 stimulation, bath application of Gly^1^-SIFamide (5 x 10^-6^ M) elicits an increase in pyloric cycle period (Blitz et al., 2019). Thus, the influence of the pyloric pacemaker ensemble via electrical coupling to LPG is likely similar in the two conditions. However, unlike the MCN5 inhibition of LP (Norris et al., 1996) (Fig 2A), Gly^1^-SIFamide application excites LP (Blitz et al., 2019) (Fig 2B). The previous characterization of MCN5 inhibition of LP used brief (10 s) MCN5 stimulation. To better compare the effects of MCN5 and its bath applied transmitter on LP, we measured LP activity during sustained tonic MCN5 stimulation and Gly^1^-SIFamide application. We found that LP activity during Gly^1^-SIFamide application was stronger in terms of number of spikes per burst and firing rate compared to during MCN5 stimulation (LP # spikes/burst: MCN5: 3.51 ± 0.70; Gly^1^- SIFamide: 7.14 ± 0.61; t(36) = -3.42, p = 0.002; LP spike frequency: MCN5: 11.46 ± 1.76 Hz; Gly^1^-SIFamide: 18.31 ± 1.45 Hz; t(36) = -2.70, p = 0.011; Student’s t-test, MCN5, n =11; Gly^1^-SIFamide, n = 27).

Since LP inhibits LPG, we explored whether stronger LP activity during Gly^1^- SIFamide application prevented LPG from generating gastric mill-timed (slow) bursts that occur during MCN5 stimulation. We first examined whether there was a relationship between LP activity and LPG slow bursting. In a plot of LP firing frequency versus LP number of spikes per burst, with the number of LPG slow bursts color-coded, it is evident that at lower LP number of spikes and firing rate, there were more slow bursts (warmer colors) for both MCN5 stimulations (filled circles) and Gly^1^-SIFamide applications (open circles) (Fig. 3A). At higher LP firing frequencies and numbers of spikes per burst, there were fewer LPG slow bursts, evident in the cooler colors (Fig. 3A). We found that across the range of LP activity measured, there was a negative correlation between number of LPG slow bursts and LP number of spikes per burst and spike frequency in Gly^1^-SIFamide (Fig. 3A; # LPG slow bursts vs LP # of spikes/burst: r^2^= -0.58, p = 0.002; # LPG slow bursts vs LP spike frequency, r^2^ = -0.62, p = 8.5 x 10^-4^, Pearson Correlation; n = 25). Although average LP activity was lower during MCN5 stimulation compared to Gly^1^-SIFamide, there was also a negative correlation between number of LPG slow bursts and LP activity during tonic MCN5 stimulation (Fig. 3A; # LPG slow bursts vs LP number of spikes/burst: r^2^ = -0.62, p = 0.042; #LPG slow bursts vs LP spike frequency, r^2^ = -0.60, p = 0.049; Pearson Correlation; n = 11). These data support stronger LP activity in Gly^1^-SIFamide application as a cause of the lower LPG dual-network activity in Gly^1^-SIFamide compared to during MCN5 stimulation. Before we further tested that hypothesis, we first addressed the different effects of Gly^1^-SIFamide and MCN5 on LP. Specifically, we asked whether MCN5 uses a co-transmitter to inhibit LP activity.

**Figure 3.**
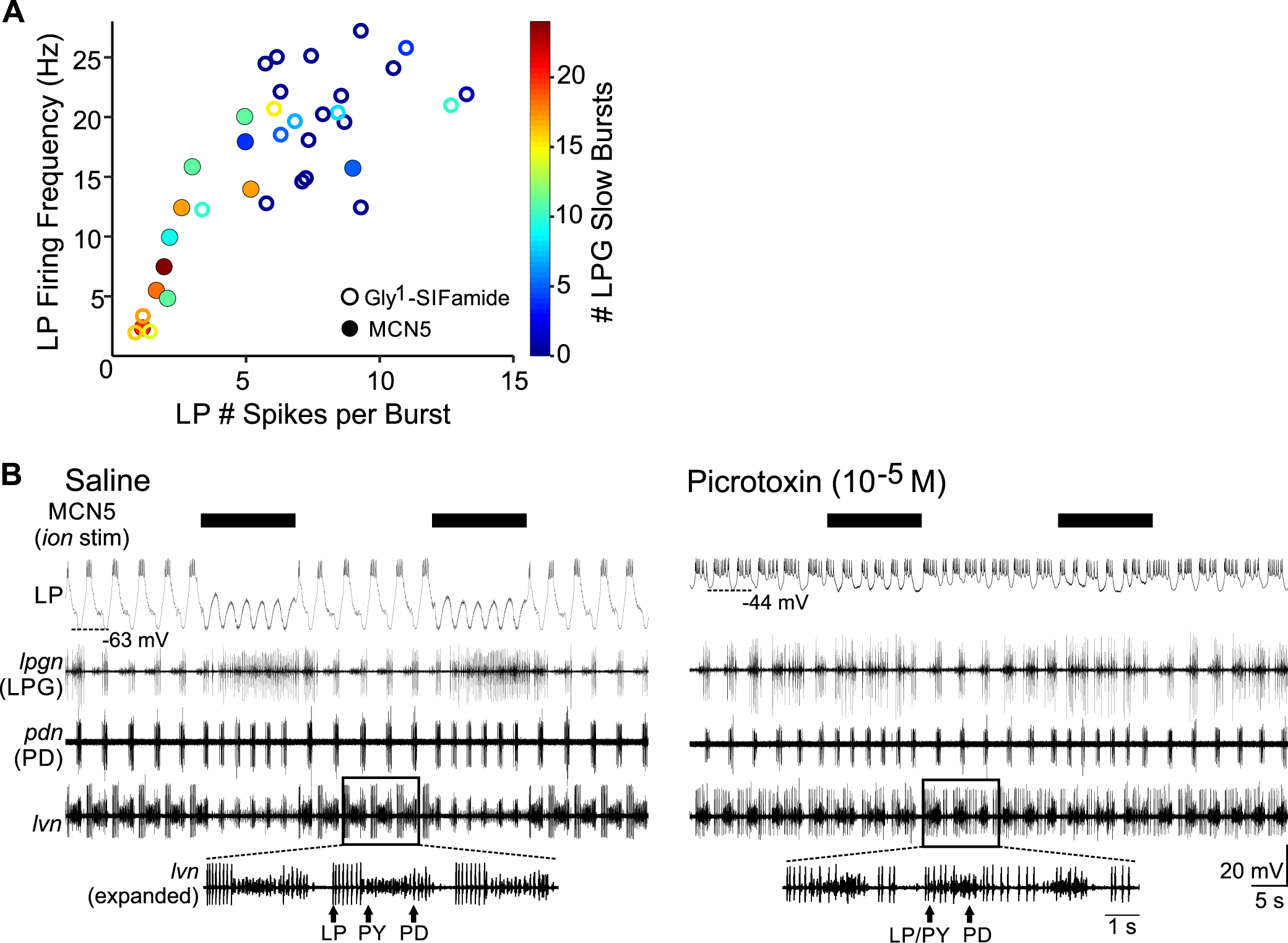
The number of Gly^1^-SIFamide- and MCN5-elicited LPG slow bursts negatively correlates with LP activity. *A*, A plot of LP firing frequency and LP spikes per burst indicates that during MCN5 stimulation (filled circles) and Gly^1^-SIFamide application (open circles), there is a greater number of LPG slow bursts (warmer colors) with weaker LP activity. Note that MCN5 data points mostly occur toward the left of the graph due to weaker LP activity and Gly^1^-SIFamide data points occur mostly toward the upper right due to stronger LP activity. In several experiments, Gly^1^-SIFamide elicited 0 LPG slow bursts (dark blue open circles). MCN5: n = 11; Gly^1^-SIFamide: n = 25. B, (*Left*) Rhythmic MCN5 stimulation (30 Hz, 5 s; black bars) inhibited LP evident by the elimination of spiking and the more hyperpolarized peak membrane potential. During each MCN5 stimulation, there was a slow burst elicited in LPG (*lpgn*). The expanded region of the extracellular lvn recording highlights that the LP and PY neurons fire out of phase, due to their reciprocal inhibition which has a stronger effect than the electrical coupling between them. (*Right*) In the presence of picrotoxin (PTX: 10^-5^ M), MCN5 stimulation increased the pyloric frequency and elicited slow bursts in LPG, however, it did not inhibit LP. Baseline LP activity is different in PTX due to the block of glutamatergic inhibition, including that from the AB/LPG and PY neurons (Bidaut, 1980; Marder and Eisen, 1984; Graubard and Hartline, 1987; Cleland and Selverston, 1995; Mamiya et al., 2003). The efficacy of PTX is evident in the expanded lvn trace in which LP and PY now overlap because glutamatergic inhibition between them is blocked, but their electrical coupling remains. LP still bursts in pyloric time due to the remaining inhibition from the cholinergic PD neuron. (Left) and (Right) are from the same preparation.

A common inhibitory neurotransmitter in the STNS is glutamate (Marder and Eisen, 1984; Cleland and Selverston, 1995), thus we tested whether MCN5 uses glutamate to inhibit LP. Rhythmic MCN5 stimulation (intra-burst frequency: 30 Hz; inter-burst frequency: 0.1 Hz, burst duration: 5 s) was used to assess MCN5 inhibition of LP. In control saline conditions, LP was bursting in pyloric time (6.0 ± 0.9 spikes/burst; firing frequency: 15.5 ± 2.1 Hz) with a voltage at the peak of its oscillations of -40.6 ± 2.0 mV (Fig. 3B, left) (n = 6). During MCN5 stimulation, LP peak voltage during each oscillation was more hyperpolarized, and LP spike number and spike frequency decreased (Fig. 3B, left) (LP peak voltage: -49.2 ± 2.8 mV, t(5) = 4.40, p = 0.007; LP # spikes/burst: 2.3± 0.7, t(5) = 3.8, p = 0.01, Paired t-test; LP spike frequency (Hz): 7.3 ± 2.7, Z-Statistic =-2.2, p = 0.031, Wilcoxon Signed Rank Test; n = 6). Also note that LPG generated a gastric mill-timed burst during each MCN5 stimulation (Fig. 3B, *lpgn*). To test whether MCN5 inhibition of LP was mediated by glutamate, we used picrotoxin (PTX, 10^-5^ M), which blocks inhibitory glutamatergic transmission in the STNS (Bidaut, 1980; Marder and Eisen, 1984; Cleland and Selverston, 1995). To verify that PTX effectively blocked glutamatergic inhibition, we used LP and PY neuron activity. These neurons are electrically coupled and reciprocally inhibit each other via glutamatergic synapses (Graubard and Hartline, 1987; Mamiya et al., 2003). Under baseline conditions, the chemical inhibition dominates and LP and PY activity is largely non-overlapping (Mamiya et al., 2003) (Fig. 3B, *left expanded lvn*). Blocking glutamatergic inhibition with PTX eliminates LP and PY chemical inhibition of each other, leaving only electrical coupling, such that their activity is coincident (Bidaut, 1980). Thus, overlapping LP and PY activity was used to verify the efficacy of PTX (Fig. 3B, right expanded lvn). In PTX, LP peak voltage was less hyperpolarized at -35.2 ± 0.8 mV and its burst was extended due to the block of PY inhibition, resulting in 10.3 ± 0.7 spikes per burst, and a spike frequency of 11.5 ± 0.7 Hz (Fig. 3B, right). In this condition, MCN5 stimulation decreased LP spike number (6.2 ± 0.8, t(5) = 5.44, p = 0.003, Paired t-test, n = 6), but not spike frequency (11.0 ± 0.7 Hz, t(5) = 1.49, p = 0.197, Paired t-test, n = 6) or peak voltage (-35.6 ± 0.9 mV, t(5) = 2.25, p = 0.074, Paired t-test, n = 6) (Fig. 3B, right). In PTX, the decreased LP number of spikes per burst caused by MCN5 stimulation was likely due to PD neuron inhibition of LP, as the PD neurons are cholinergic, and their chemical inhibitory transmission is not blocked by PTX (Marder and Eisen, 1984). The increased PD burst frequency during MCN5 stimulation, truncated the LP burst duration (Fig. 3B, right) (PTX: 0.8 ± 0.07 s; PTX + MCN5 stimulation: 0.5 ± 0.07 s; t(5) = 3.8, p = 0.013, Paired t-test, n = 6), resulting in fewer LP spikes per burst. However, the lack of a change in LP spike frequency and peak voltage in response to MCN5 stimulation indicates that the direct MCN5 inhibition of LP was blocked by PTX. Thus, MCN5 appears to use glutamate as a co-transmitter to inhibit LP, whereas its modulation of LPG occurs via Gly^1^-SIFamide (Fig. 3) (Blitz et al., 2019). There are two Gly^1^-SIFamide-containing inputs projecting from each CoG to the STG (Blitz et al., 2019) and thus the excitation of LP likely mimics the actions of the other, unidentified Gly^1^-SIFamide modulatory neuron. The negative correlations between LP activity and LPG slow bursts coupled with the distinct transmitters used by MCN5 to modulate LPG and inhibit LP support our hypothesis that the Gly^1^-SIFamide excitation of LP interferes with its ability to elicit dual bursting activity in LPG.

Although the number of LPG slow bursts is useful to determine LPG participation in the gastric mill rhythm, it does not provide a measure of LPG dual-network activity.

Thus, to quantify LPG dual-network activity, we used spectral analysis to determine the extent to which LPG participates in the pyloric and gastric mill rhythms (Fig. 4) (Bucher et al., 2006; Rehm et al., 2008) (see Methods). Briefly, spectral analysis quantifies the extent of spiking activity across frequencies (Bucher et al., 2006; Rehm et al., 2008).

**Figure 4.**
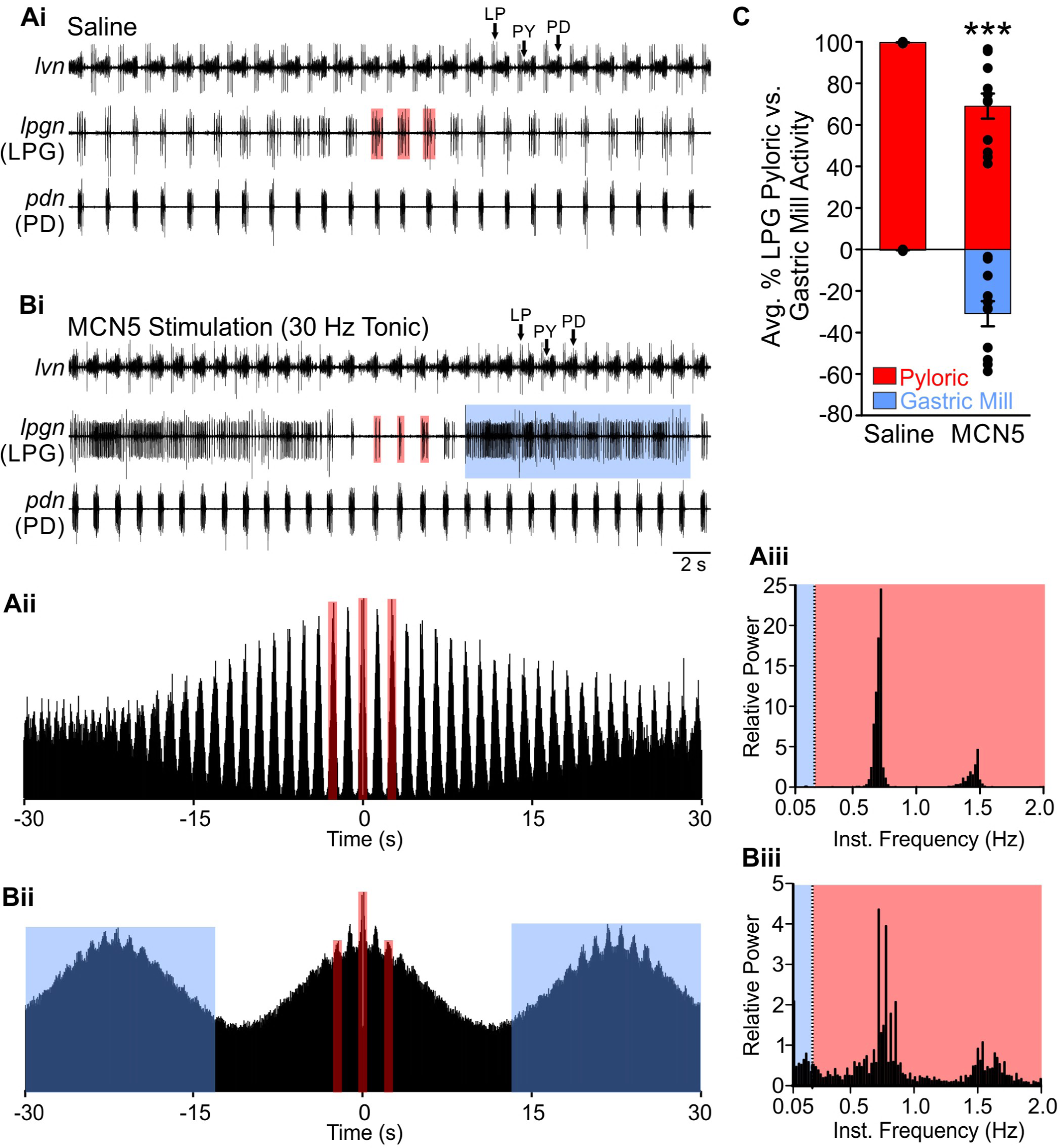
Spectral analysis enables quantification of LPG dual-network activity. Ai, Extracellular recordings of the *lvn* (LP, PY, PD), *lpgn* (LPG), and *pdn* (PD) illustrate pyloric only activity, including in LPG (red boxes) in control conditions. *Aii*, An autocorrelogram of (100 s) of LPG activity in this experiment results in narrow peaks that identify the pyloric-timed bursts. For illustrative purposes, autocorrelogram plots are truncated at ± 30 s. *Aiii*, To quantify the pyloric vs gastric mill timed activity, the relative power across a range of instantaneous frequencies (IF; see Methods) was plotted. The blue area on the graph highlights the gastric mill interburst frequencies (0.05 – 0.2 Hz), and the red area highlights the pyloric interburst frequencies (0.2 – 4.0 Hz). In control, the spectral analysis indicates almost entirely pyloric-timed activity. For illustrative purposes, histogram plots are truncated at 2.0 Hz. Bi, During MCN5 stimulation, LPG dual activity is evident in the extracellular recording. LPG dual-network activity is also evident as broad peaks (blue boxes) and narrower peaks (red boxes) in the autocorrelogram (*Bii*) and as peaks of relative power occurring within gastric mill (blue region) and pyloric (red regions) frequencies in the relative power histogram (*Biii*). The total power from 0.05 - 4.0 Hz was summed across such a histogram for each condition, in each experiment, and the gastric mill and pyloric components calculated as a percentage of the total. These percentages were then averaged across experiments, with pyloric plotted as a positive value and gastric mill plotted as a negative value for illustrative purposes. C, Across experiments, a larger percentage of LPG activity was gastric mill-timed (blue box) during MCN5 stimulation compared to control. Dots represent individual experiments, in control the values were very close to 100% and 0% for pyloric and gastric mill-timed, respectively, and thus dots overlap. Mean ± SE; n = 11 each condition, p = 5.1 x 10^-4^, paired t-test. (A) and (B) are from the same preparation.

The approach we took is illustrated in figure 4. In control, the LPG neuron generated bursts in pyloric time, coincident with the PD neuron (Fig. 4Ai, red boxes). An autocorrelogram graphically depicts the LPG pyloric-timed bursting pattern by plotting the timing of each LPG action potential relative to every other LPG action potential across a stretch of time (Bucher et al., 2006), such that each peak corresponds to an LPG pyloric-timed burst (Fig. 4Aii, red boxes). We then generated a histogram of the relative power of LPG activity across a range that included inter-burst spike frequencies (0.05 - 4 Hz) but not intra-burst spike frequencies. From the histogram, the relative power was summed across inter-burst spike frequencies corresponding to gastric mill timing (0.05 - 0.19 Hz;) (Fig. 4Aiii, blue area) and pyloric timing (0.2 - 4.0 Hz) (Fig. 4Aiii, red area, x-axis truncated at 2.0 Hz in the example for viewing ease). These values were normalized to the total sum of relative power across gastric mill and pyloric-timed inter-burst spike frequencies, converted to a percentage, and averaged across experiments (Fig. 4C) (see Methods). Gastric mill percent values are graphically represented as negative values. In control in the example shown (Fig. 4A), the percent of LPG activity corresponding to gastric mill and pyloric timing was 0.5 % and 99.5 %, respectively. During tonic (30 Hz) MCN5 stimulation, LPG burst in both pyloric and gastric mill time (Fig. 4Bi, red and blue boxes, respectively). Note that during MCN5 stimulation LP fired 0-2 spikes per burst, whereas it fired 4-6 spikes per burst in the saline condition. The autocorrelogram of LPG activity during MCN5 tonic stimulation has two peaks with two different sets of characteristics: 1) tall, narrow peaks associated with LPG pyloric-timed bursts, similar to control (Fig. 4Aii; and 2) broad peaks associated with LPG gastric mill-timed bursts during tonic MCN5 stimulation (Fig. 4Bii, red and blue boxes, respectively). The histogram of the relative power of LPG bursting activity during MCN5 tonic stimulation revealed activity in both gastric mill and pyloric frequencies, with LPG gastric mill and pyloric percentages of 47.3 % and 52.7 %, respectively (Fig. 4Biii). Using this method of quantifying LPG bursting, the percent of gastric mill timed LPG activity was greater during tonic MCN5 stimulation compared to control (Fig. 4C; Control: 0.3 ± 0.07; MCN5: 31.0 ± 6.1; t(10) = -5.04, p = 0.001, Paired t-test, n = 11).

We next utilized spectral analysis to address whether LP activity was responsible for the difference in LPG activity between MCN5 and Gly^1^-SIFamide application. In particular, we tested whether LP hyperpolarization during Gly^1^-SIFamide application would enable LPG to switch to dual-network activity, similar to its response to MCN5 stimulation. In Gly^1^-SIFamide (5 x 10^-6^ M), LP was active, and LPG generated bursts in pyloric time, with some bursts extending beyond the end of a PD burst (Fig. 5A, left traces, red boxes) (Blitz et al., 2019). Similar to the analysis involving number of LPG gastric mill-timed bursts, the percentage of LPG gastric mill-timed activity calculated via spectral analysis differed between Gly^1^-SIFamide application and tonic MCN5 stimulation (Fig. 5B) (MCN5: 31.0 ± 6.1 %, n = 11; Gly^1^-SIFamide: 10.1 ± 3.6 %, n = 15; H(3) = 20.93, p = 0.02, Kruskal-Wallis One-way ANOVA on Ranks, Dunn’s post hoc).

**Figure 5.**
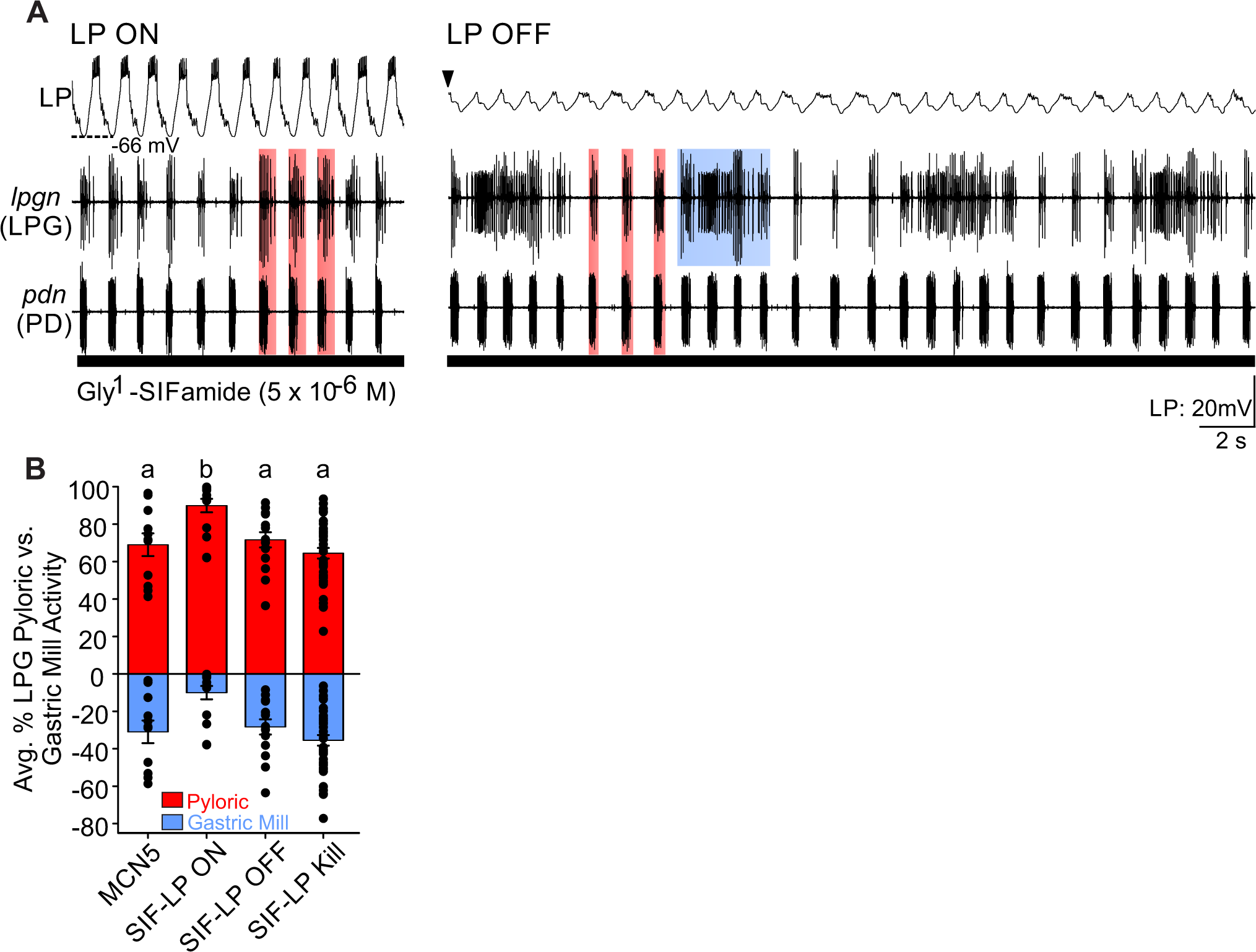
Gly^1^-SIFamide more closely mimics MCN5-elicited effects in the absence of LP. A, (*Left*) With LP intact, LPG is primarily pyloric-timed in Gly^1^-SIFamide (5 x 10^-6^ M) with periodic (*lpgn*) bursts that extend beyond the end of the pyloric PD bursts (red boxes), but no slow gastric mill-timed bursts. (*Right*) During the Gly^1^-SIFamide application, hyperpolarization of LP caused LPG bursting to switch to both pyloric (red boxes) and gastric mill (blue boxes) timed. B, Across experiments, the average percent of LPG pyloric (red) and gastric mill (blue) activity was similar between MCN5 stimulation, Gly^1^-SIFamide with LP hyperpolarized (SIF-LP OFF) and Gly^1^-SIFamide with LP photoinactivated (SIF-LP Kill; see Methods). However, the LPG activity was less gastric-mill timed during Gly^1^-SIFamide application with LP active (SIF-LP ON). MCN5 (n=11), SIF-LP ON (n = 15), SIF-LP OFF (n = 15), and SIF-LP Kill (n = 39). Similar letters indicate no statistical difference, different letters indicate significance, Kruskal-Wallis One Way ANOVA on Ranks, Dunn’s post-hoc.

However, when LP was hyperpolarized during Gly^1^-SIFamide, the percentage of LPG gastric mill-timed activity increased, becoming similar to that during tonic MCN5 stimulation (Fig. 5A-B: Gly^1^-SIFamide-LP OFF: 28.4 ± 4.1, H(3) = 20.93, p = 1.0, Kruskal-Wallis One-way ANOVA on Ranks, Dunn’s post hoc). Although hyperpolarizing LP enabled LPG to switch to dual-network activity in Gly^1^-SIFamide, it was possible that the current injection into LP was influencing other network neurons via electrical coupling (Fig. 1B) (Graubard and Hartline, 1987; Mamiya et al., 2003; Marder and Bucher, 2007). We thus moved to photoinactivation, which enables the permanent removal of a neuron without affecting electrically coupled neurons (Miller and Selverston, 1979; Marder and Eisen, 1984; Faumont et al., 2005) (see Methods). After photoinactivation (“Kill”) of LP, LPG generated dual-network activity in Gly^1^-SIFamide that was similar to that during Gly^1^-SIFamide with LP hyperpolarized, and to tonic MCN5 stimulation (Fig. 5B; Gly^1^-SIFamide-LP Kill vs. Gly^1^-SIFamide-LP OFF: H(3) = 20.93, p > 0.05; Gly^1^-SIFamide-LP Kill vs. MCN5 Tonic: H(3) = 20.93, p > 0.05; SIF-LP Kill, n = 39; SIF-LP ON, n = 15; MCN5 tonic, n = 11; Kruskal-Wallis One-way ANOVA on Ranks, Dunn’s post hoc). These results indicate that Gly^1^-SIFamide application with LP activity eliminated more closely mimics MCN5-elicited actions and enables us to examine mechanisms underlying LPG switching to dual-network activity. Therefore, for the remainder of this study, Gly^1^-SIFamide applications were performed with the LP neuron photoinactivated unless otherwise noted.

Gastric mill neuron input is not necessary for LPG to generate dual-network activity

Strengthening chemical synaptic input from network neurons to recruit a switching neuron into another network is the established mechanism for neuronal switching (Dickinson et al., 1990; Hooper and Moulins, 1990; Meyrand et al., 1994; Weimann and Marder, 1994). Therefore, we tested whether LPG required synaptic input from gastric mill neurons to switch to dual-network activity. There are eight neurons that can participate in a gastric mill rhythm (Fig. 1B) (Weimann et al., 1991). In other gastric mill rhythm versions, the reciprocally inhibitory LG and interneuron 1 (Int1) neurons are the core rhythm generators (Coleman et al., 1995; White and Nusbaum, 2011). LG changes from silent to bursting upon activation of a gastric mill rhythm, including the MCN5/Gly^1^-SIFamide gastric mill rhythm (Coleman et al., 1995; Beenhakker et al., 2004; White and Nusbaum, 2011; Blitz et al., 2019). In the absence of a gastric mill rhythm, Int1 is active in pyloric time due to rhythmic inhibition from the pyloric pacemakers (Fig. 1B) (Weimann et al., 1991; Bartos et al., 1999). During a gastric mill rhythm however, its pyloric-timed activity is rhythmically inhibited in gastric mill time by LG (Coleman et al., 1995; Bartos and Nusbaum, 1997; White and Nusbaum, 2011).

Using Int1 inhibitory postsynaptic potentials (IPSPs) in LG, we verified that Int1 is active with both the pyloric and gastric mill rhythm during Gly^1^-SIFamide application (n = 11/11). To determine whether LG and Int1 are core members of the Gly^1^-SIFamide rhythm generator and thus necessary for LPG slow bursting, we eliminated LG activity with hyperpolarizing current injection, which also changed Int1 activity to pyloric only (n = 11/11). As is evident in an example experiment, LPG slow bursting continued during Gly^1^-SIFamide in the absence of alternating LG/Int1 activity (Fig. 6A, right), indicating that LPG dual activity does not require gastric mill-timed input from these two neurons. This result also suggests that the Gly^1^-SIFamide gastric mill rhythm relies on a rhythm generator that is different from other gastric mill rhythms (Coleman et al., 1995; White and Nusbaum, 2011).

**Figure 6.**
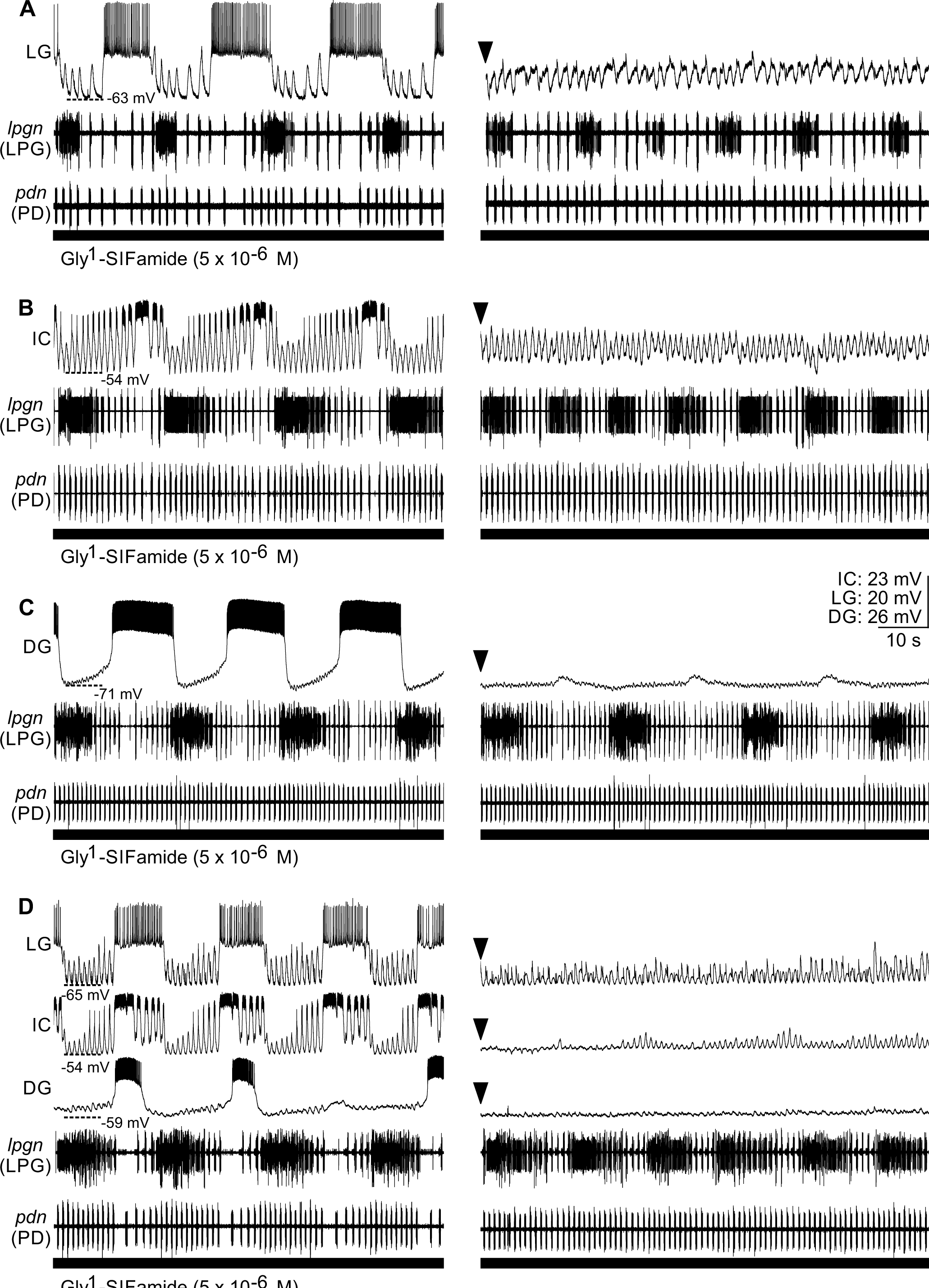
The LPG switch to dual bursting does not require activity from other gastric mill network neurons during Gly^1^-SIFamide bath application. *A-D*, (*Left*) Gly^1^-SIFamide application with LP photoinactivated elicited dual-network bursting in LPG (*lpgn*). (*Right*) LPG continued to express dual bursting when LG (*A*), IC (*B*), DG (*C*), or LG, IC, and DG (D) activity was eliminated with hyperpolarizing current injection (*right, downward-pointing arrowheads*). During current injections, neurons were recorded with an unbalanced bridge and hyperpolarized traces are aligned for ease of visualization, not to represent actual membrane potentials. A-D are from separate experiments.

In the MCN5/Gly^1^-SIFamide gastric mill rhythm, in addition to LG, the IC and DG neurons are most strongly activated in gastric mill time (Blitz et al., 2019). Thus, we examined whether these neurons are necessary to recruit LPG into the gastric mill network and generate dual-network activity. We injected hyperpolarizing current in each neuron, individually (Figs. 6B-C) and simultaneously (Fig. 6D), to remove their activity during Gly^1^-SIFamide (5 x 10^-6^ M) bath application. LPG continued to generate slower gastric-mill timed bursts and thus dual-network activity during hyperpolarization of the IC neuron (Fig. 6B, right, DG neuron (Fig. 6C, right), and LG, IC, DG neurons, simultaneously (Fig. 6D, right). However, there appeared to be differences in some aspects of LPG gastric mill-timed bursting, and pyloric rhythm regularity during IC hyperpolarization (Fig. 6B, right). This is consistent with the identified correlation between IC burst duration and pyloric cycle period during Gly^1^-SIFamide application and MCN5 stimulation (Blitz et al., 2019), and IC inhibition of the pyloric pacemakers during activation of modulatory commissural neuron 7 (MCN7; Blitz et al. 1999), but we did not explore it further here. To quantify the LPG dual-network activity, we again used spectral analysis (Table 1, Fig. 7). As in the examples shown, across preparations, LPG activity had a gastric component that was similar to before and after the neuron was hyperpolarized for IC and DG (Fig. 7B-C, blue bars; Table 1). Although the percentage of LPG dual-network activity was statistically similar to saline conditions during LG hyperpolarization (Fig. 7A; Table 1), we observed LPG gastric mill-timed bursting during LG hyperpolarization (Fig. 6A). In fact, using our LPG gastric mill-timed burst criteria (see Methods), LPG generated slow bursts during LG hyperpolarization in all but one preparation (n = 16/17). Further, the number of LPG slow bursts with LG off, was not different from the number of LPG slow bursts with LG on (LG ON-Pre to OFF to ON-Post: 14.4 ± 1.1 to 15.5 ± 1.8 to 14.1 ± 1.1; F(16,2) = 0.81; ON-Pre vs OFF, p = 0.587;

**Figure 7.**
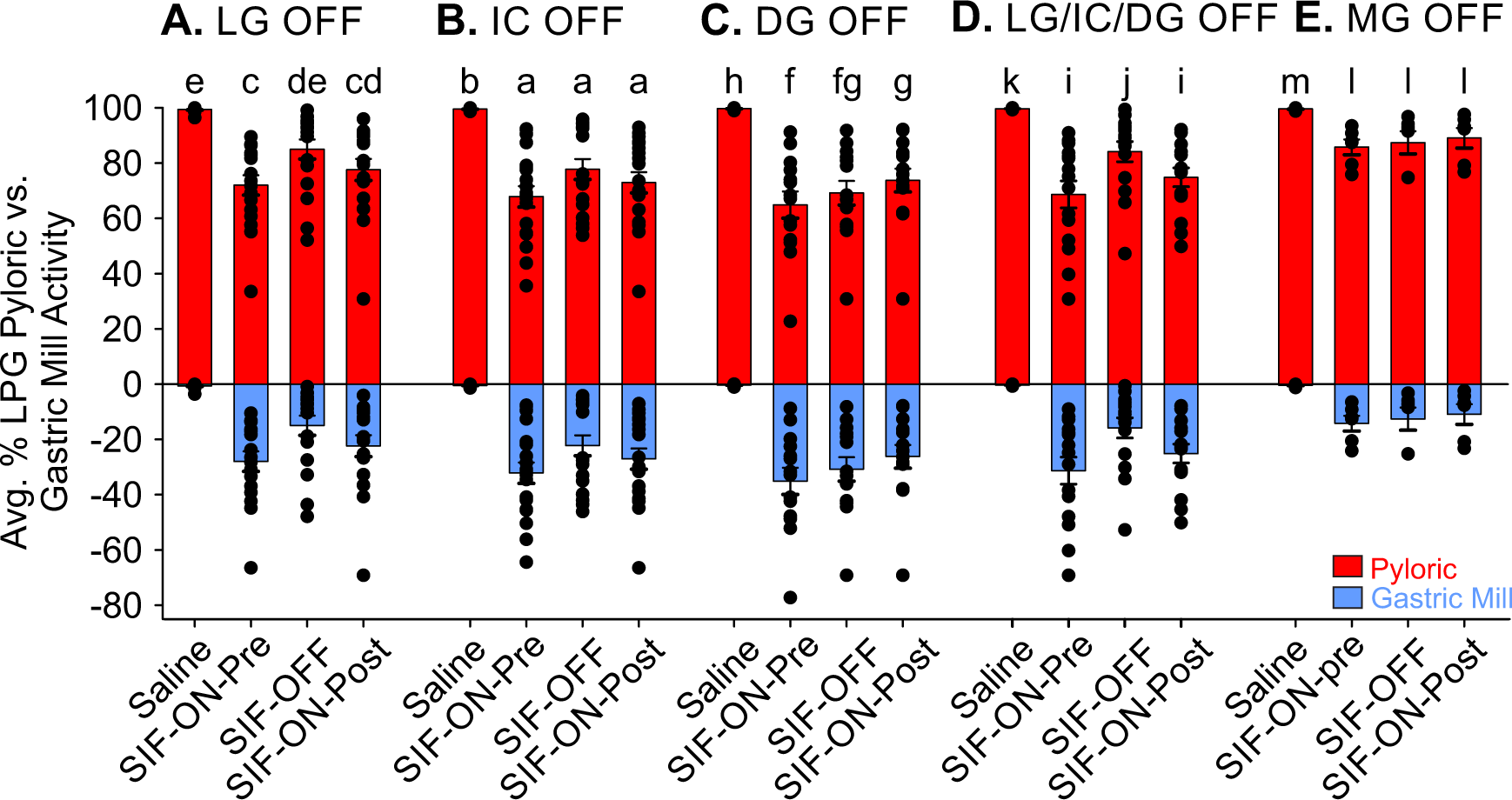
The LPG switch to dual bursting does not require activity from other gastric mill network neurons. Spectral analysis was performed to quantify the extent of pyloric and gastric mill-timed LPG activity in saline and during Gly^1^-SIFamide application before (SIF-ON-Pre), during (SIF-OFF), and after (SIF-ON-Post) hyperpolarization of LG (*A*; n = 17), IC (*B*; n = 19), DG (*C*; n = 14), LG, IC, and DG (*D*; n = 15), and MG (*E*; n = 6). (*A-C*) Friedman Repeated Measures ANOVA On Ranks, Tukey’s post hoc; (*D, E*) One-Way Repeated Measures ANOVA, Holm-Sidak post hoc. Similar letters indicate no statistical difference within each group, different letters indicate significance (see Table 1 for p-values).

**Table 1.**
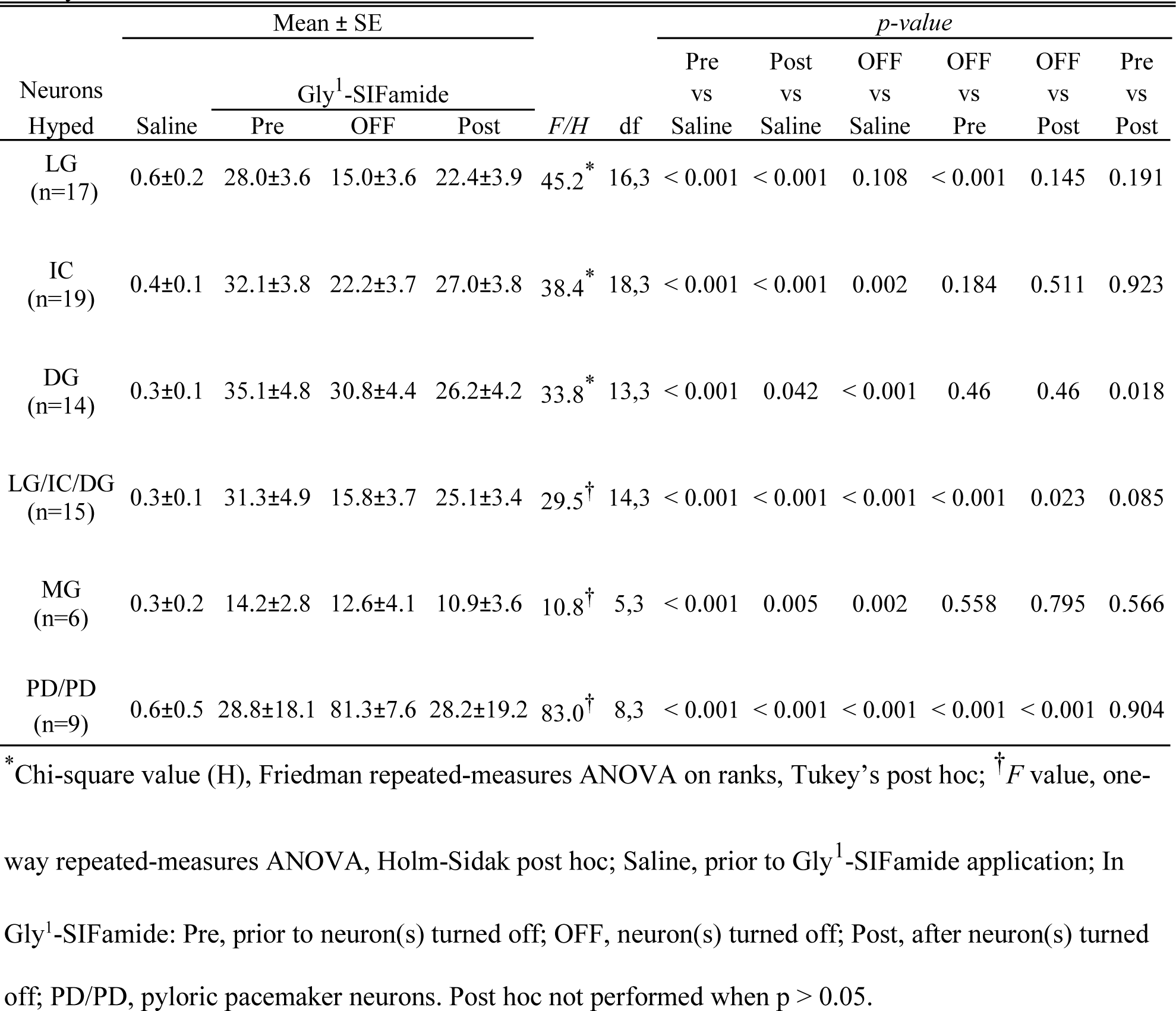
Percent of LPG gastric mill-timed bursting with/without gastric mill network and pyloric pacemaker neuron activity

ON-Post vs OFF, p = 0.630; Saline vs OFF, p < 0.001; One-way repeated measures ANOVA on ranks, Tukey’s post hoc, n = 17). This suggests that LG may regulate the strength of LPG bursting, as indicated in the spectral analysis, but is not necessary for LPG to burst in gastric mill time. Specifically, LPG slow bursting was a smaller percentage of the total activity when LG was hyperpolarized compared to pre-hyperpolarization and when LG/IC/DG were hyperpolarized compared to pre- and post-hyperpolarization (Fig. 7A, D; Table 1). This suggests that gastric mill neurons regulate aspects of the LPG dual-network activity. However, LG, IC, and DG neuron activity is not necessary for LPG dual-network activity to occur in Gly^1^-SIFamide.

Although the MG neuron is electrically coupled to the LG neuron, and thus likely hyperpolarized along with LG, we separately tested whether MG was necessary for the switch in LPG activity. We found that LPG generated dual-network activity in Gly^1^- SIFamide during MG hyperpolarization (Fig. 7E; Table 1). Among the remaining gastric mill neurons, the ventricular dilator (VD) neuron gastric mill activity is due to rhythmic inhibition from LG (Weimann and Marder, 1994) and thus VD was only active in pyloric time during hyperpolarization of LG (data not shown). As indicated above, this did not prevent dual activity in LPG. Gastric mill (GM) neurons are not excited by Gly^1^- SIFamide (Blitz et al., 2019), and thus were not examined in this study. Thus, LPG is not recruited into a second network via synaptic input from gastric mill neurons.

Pyloric pacemaker activity is not required for LPG gastric mill-timed bursting

Despite LPG not requiring gastric mill input to generate dual-network activity, it is possible that pyloric network neurons contribute to LPG switching to dual-network activity. Pyloric network input to LPG includes inhibition from LP, which was photoinactivated in our experiments, and electrical coupling to the other pacemaker ensemble neurons, the two PD and single AB neurons (Fig. 1B) (Marder and Bucher, 2007). In control conditions, AB is the pacemaker, and its rhythmic activity drives rhythmic PD and LPG bursting via electrical coupling, from AB to PD and LPG, as well as between PD and LPG (Marder and Eisen, 1984; Shruti et al., 2014; Zhang, Fahoum, Blitz, and Nusbaum, unpublished). To test for a role of the pyloric network, we applied Gly^1^-SIFamide and injected hyperpolarizing current into each of the PD neurons to turn off pyloric pacemaker activity (Fig. 8). Due to the electrical coupling, hyperpolarization of the PDs sufficiently hyperpolarizes AB, to eliminate rhythmic activity in all five pacemaker ensemble neurons (Bartos et al., 1999; Blitz et al., 2008). Shutting off rhythmic electrical synaptic input to LPG eliminates its pyloric timed bursting but does not alter its most hyperpolarized membrane potential (see Methods).

**Figure 8.**
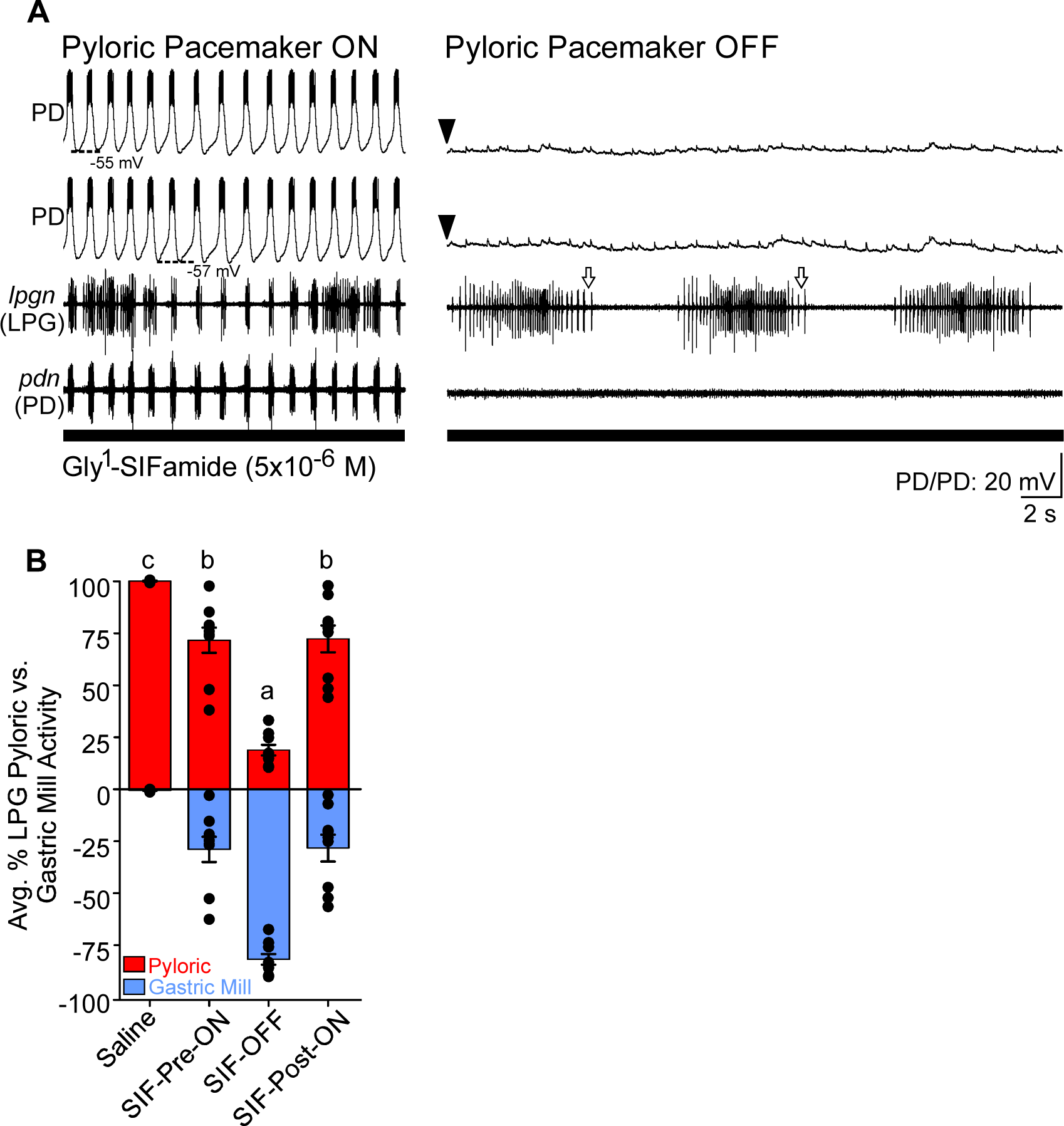
The Gly^1^-SIFamide-elicited LPG switch to gastric mill-timed bursting does not require rhythmic input from the pyloric pacemaker ensemble. *A*, (*Left*) Gly^1^-SIFamide (5 x 10^-6^ M) bath application elicited dual-network activity in LPG (*lpgn*) with the pyloric pacemaker neurons (AB, PD, PD) active. (*Right*) During the Gly^1^-SIFamide application, each of the two PD neurons was hyperpolarized (-4 nA; downward arrowheads), which also hyperpolarized the AB neuron (not shown; see Methods) and eliminated rhythmic pyloric-timed activity in the entire pacemaker ensemble, including LPG. However, LPG continued to produce slow, gastric mill-timed bursting. White arrows indicate regions of LPG slow bursting with relatively longer interspike intervals that are identified as pyloric-timed activity with our spectral analysis (see Methods) during PD hyperpolarization (*B*, SIF-OFF). Neurons were recorded with an unbalanced bridge and hyperpolarized traces are aligned for ease of visualization, not to represent actual membrane potentials. *B*, LPG activity was analyzed with spectral analysis and converted to a percentage pyloric- (red) and gastric mill- (blue) timed activity. There was more gastric mill-timed activity during all Gly^1^-SIFamide application compared to saline. Additionally, during PD neuron hyperpolarization and elimination of the rhythmic pyloric pacemaker activity (SIF-OFF), there was more gastric mill-timed LPG activity compared to before (SIF-ON-Pre) and after (SIF-ON-Post) PD hyperpolarization. Similar letters indicate no statistical difference, different letters indicate significance. One-Way Repeated Measures ANOVA, Holm-Sidak post hoc (see Table 1 for p-values).

Rhythmic pyloric network input was also not necessary for LPG to generate oscillations at a gastric mill frequency. We first verified in each experiment that Gly^1^- SIFamide (5 x 10^-6^ M) application elicited dual bursting in LPG with the pyloric rhythm on (Fig. 8A, left). We then hyperpolarized the two PD neurons to turn off the pyloric rhythm, eliminating pyloric bursting in LPG. However, LPG continued to generate gastric mill-timed bursts (Fig. 8A, right). Across experiments, in the absence of pyloric pacemaker activity, LPG generated gastric mill-timed bursting but failed to produce pyloric-timed bursts (n = 9/9) (Table 1, Fig. 8B). The small amount of LPG activity that occurred within the spectral range of pyloric timing reflects longer ISIs at the margins of the gastric mill-timed bursts (e.g., white arrows, Fig. 8A, right). The inability to maintain pyloric-timed bursting indicates that AB continued to be a necessary pacemaker neuron for the pyloric rhythm and LPG was co-active with the pyloric rhythm due to electrical coupling. However, the ability of LPG to generate gastric mill-timed bursts without input from the pyloric pacemaker neurons, or the gastric mill neurons (above) suggests that LPG switches into the gastric mill pattern due to Gly^1^-SIFamide modulation of LPG intrinsic properties.

LPG Slow Bursting Occurs via Intrinsic Mechanisms

To ensure that there were no compensatory synaptic mechanisms supporting LPG bursting by one network when rhythmic input from the other one was eliminated, we tested whether the slow bursting elicited by Gly^1^-SIFamide, either exogenously applied or neuronally released by MCN5, occurred without rhythmic input from either network. In control Gly^1^-SIFamide application, LPG generated dual-network activity in pyloric and gastric mill time (Fig. 9Ai). We then isolated LPG from rhythmic gastric mill and pyloric input using picrotoxin (PTX, 10^-5^ M) application to block glutamatergic inhibitory chemical transmission plus PD/PD hyperpolarization (PTX:PDhype). In these experiments, the LP neuron remained intact, however its influence on LPG is blocked by PTX. During Gly^1^-SIFamide application, the isolated LPG still generated bursts (Fig. 9Aii). To confirm that this activity occurred at the slower, gastric mill frequency, we plotted cycle period and burst duration of isolated LPG activity and compared it to the same parameters of LPG pyloric and gastric mill-timed bursts during control Gly^1^- SIFamide application in the same experiments (n = 5). The isolated LPG bursts (grey circles) overlapped with the gastric-mill timed LPG bursts (black circles) and not the pyloric-timed bursts (white circles) (n = 5; Fig. 9Aiii).

**Figure 9.**
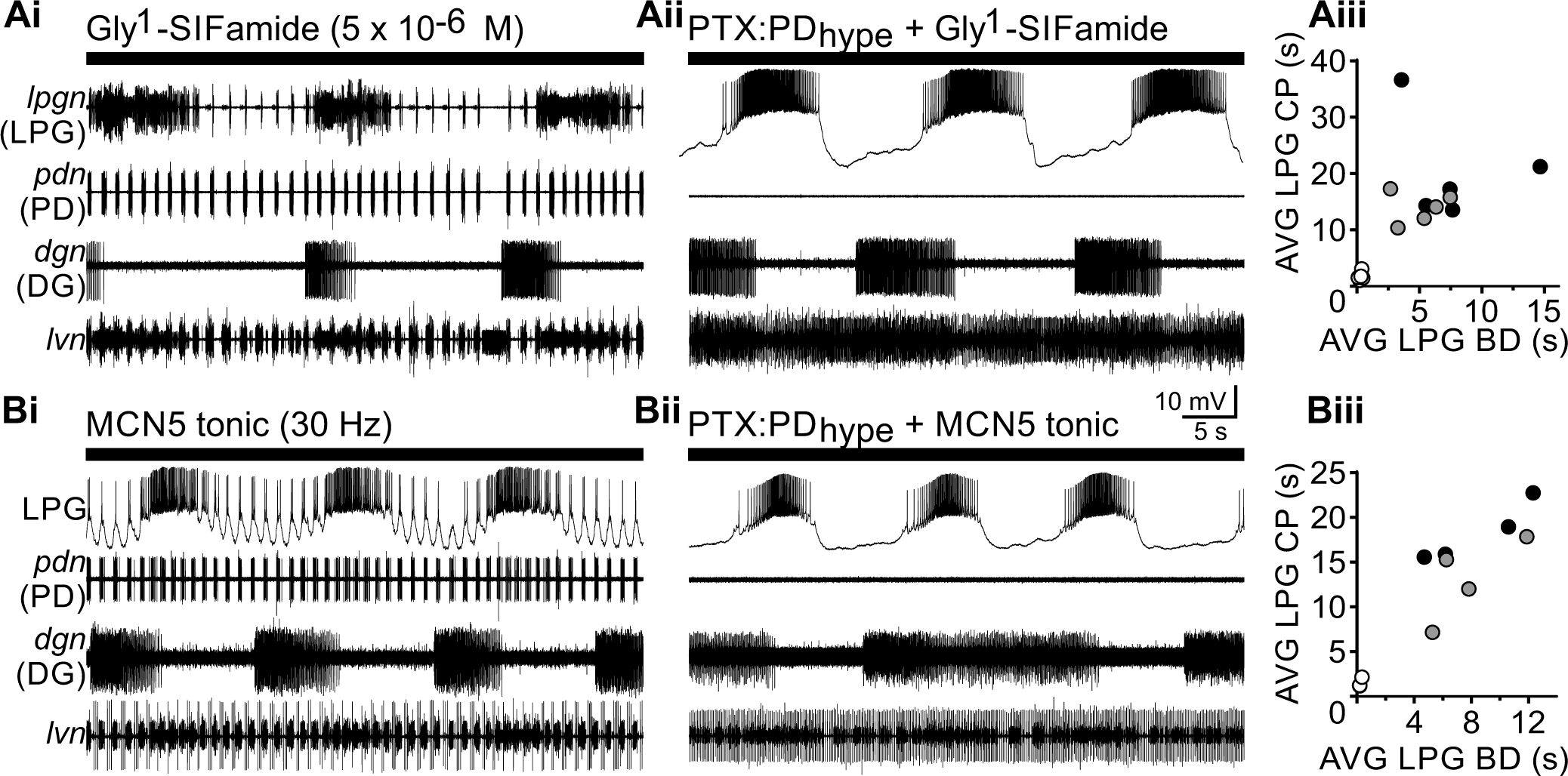
LPG gastric mill-timed bursting can be elicited by exogenous Gly^1^-SIFamide or MCN5, in the absence of any rhythmic pyloric or gastric mill network input. Ai, Bath applied Gly^1^-SIFamide (5 x 10^-6^ M) elicited rhythmic bursting in the gastric neuron DG (*dgn*), and dual bursting activity in LPG (*lpgn*). Aii, Gly^1^-SIFamide in PTX (10^-5^ M) was applied to block glutamatergic inhibition from the LP neuron and the gastric mill network, and the two PD neurons were hyperpolarized (PTX:PDhype) to eliminate rhythmic pyloric activity (*pdn*). In this isolated condition, the LPG neuron was still able to generate slow bursts. The efficacy of PTX is evident in the overlapping LP/PY activity in the lvn recording. LPG was recorded extracellularly (Ai) before an intracellular recording was obtained later in the experiment (Aii). MCN5 also elicited dual bursting in LPG during control conditions (Bi) and only slow bursting in PTX (10^-5^ M) with the two PD neurons hyperpolarized (Bii). To verify that the single frequency LPG bursting in the isolated condition was gastric mill-timed bursting, the average cycle period (CP) and burst duration (BD) was plotted for pyloric and gastric mill LPG bursts in control conditions, and for isolated LPG bursts. The isolated LPG bursts (grey circles) cluster with control condition gastric mill bursts (black circles) and not with pyloric bursts (white circles), during both Gly^1^-SIFamide application (*Aiii*) and MCN5 stimulation (*Biii*). Gly^1^- SIFamide, n = 5; MCN5, n = 4.

MCN5 stimulation also elicited slow, gastric-mill timed bursting in LPG when isolated from both networks. MCN5 stimulation (30 Hz) elicited dual-network activity in control conditions (Fig. 9Bi) and only slow bursting in the isolated condition (PTX:PDhype) (Fig. 9Bii). As with bath-applied Gly^1^-SIFamide, during MCN5 stimulation, the slow LPG bursting when isolated from both networks (grey circles) aligned with gastric mill-timed bursting (black circles) and not pyloric bursts (white circles) (n = 4; Fig, 9Biii). These data indicate that the LPG dual-network activity includes fast pyloric bursting via electrical coupling, but slower gastric mill-timed bursting via intrinsic properties. Thus, if the slower LPG bursting is intrinsically generated, it should be voltage-dependent (Hablitz and Johnston, 1981; Adams and Benson, 1985; Marder and Calabrese, 1996).

We used PTX:PDhype to isolate LPG from both networks and current injections to manipulate LPG membrane potential to examine whether Gly^1^-SIFamide (5 x 10 ^-6^ M)- elicited LPG bursting was voltage-dependent. As above, the LP neuron was intact, but its actions blocked by PTX. In an example experiment, at baseline conditions (0 nA), LPG was bursting with a cycle period of 28 s and a duty cycle of 27 % (Fig. 10A). Duty cycle is the percentage of a cycle during which a neuron is active. Depolarization of LPG (+1 nA) noticeably increased LPG burst duration, with a smaller effect on cycle period (22 s), resulting in an increased duty cycle (70 %). A further depolarization (+1.5 nA) resulted in tonic firing, although there was still some visible regulation of firing rate (Fig. 10A). Hyperpolarizing LPG (-1 nA) decreased burst duration and increased cycle period (43 s), thus decreasing duty cycle (6 %). Further hyperpolarization (-2 nA) eliminated LPG bursting (Fig. 10A). Electrical EPSPs in the LPG intracellular trace are from the other LPG neuron, as these two neurons are electrically coupled (Weimann et al., 1991). Across multiple experiments, the LPG cycle period increased with hyperpolarizing current injections and increased with depolarizing current injections (Fig. 10B). This parabolic relationship of cycle period with voltage occurs in other intrinsically bursting neurons based on the balance of inward and outward conductances (Skinner et al., 1993; Selverston et al., 2009). Bursting was eliminated (grey bars) with sufficiently hyperpolarizing current injections (-1 nA, n = 1/6; -1.5 nA, n = 1/6; -2 nA, n = 4/6) (Fig. 10B). In some experiments, LPG fired tonically (white bar) based on our burst criteria (see Methods), such that there were no ISIs longer than a typical pyloric cycle with depolarizing current injections (1.5 nA, n = 3). For three preparations in which LPG did not fire tonically, LPG was bursting at the highest current injections where the recording was still stable (1 nA, n = 1/6; 1.5 nA, n = 2/6). LPG duty cycle decreased with hyperpolarizing current injection and increased with depolarizing current injection (Fig. 10C). Collectively, these data identify LPG slow gastric mill-timed bursting elicited by the modulatory actions of bath-applied and MCN5-released Gly^1^- SIFamide as voltage-dependent, intrinsically generated.

**Figure 10.**
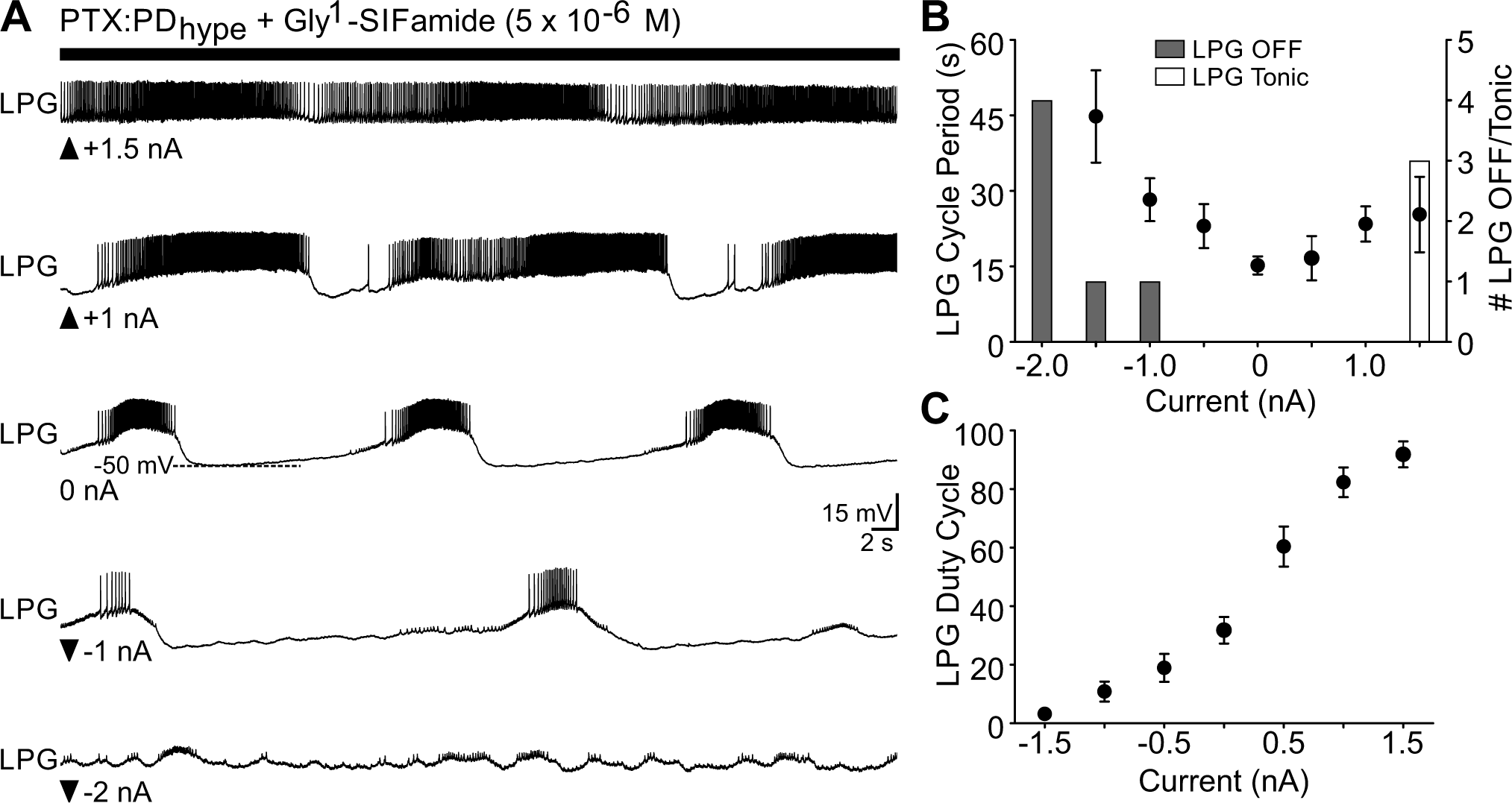
LPG gastric mill-timed bursting is voltage dependent. *A*, Several current injection levels are shown for an example LPG neuron during PTX:PDhype and Gly^1^- SIFamide application. Depolarizing current injections (upward-facing arrowheads) increased LPG burst duration (+1) or elicited tonic firing (+1.5 nA). Activity was considered to be tonic when there were no ISIs longer than a typical pyloric cycle. Hyperpolarizing current injections (downward-facing arrowheads) increased cycle period and decreased burst duration (-1 nA) or eliminated bursting (-2 nA). *B*, The average LPG cycle period across experiments is plotted as a function of current injection (circles ± error bars). The numbers of preparations in which LPG bursting turned off (grey bars) or became tonic (white bar) are plotted on the same graph (n = 6). *C*, LPG duty cycle, the percentage of a cycle that a neuron is active, is plotted as a function of current injection. LPG duty cycle decreased with hyperpolarizing current injection and increased with depolarizing current injection. Bursting parameters are the average of 2 - 5 cycles at each current injection (see Methods). (n = 6).

## Discussion

We identified a novel mechanism of switching a neuron from single-to dual-network participation. Specifically, modulating intrinsic membrane properties enables neuronal switching into bursting at a second network frequency (LPG - gastric mill rhythm), while electrical synapses maintain switching neuron activity in its “home” network (LPG - pyloric rhythm) (Fig. 11).

**Figure 11.**
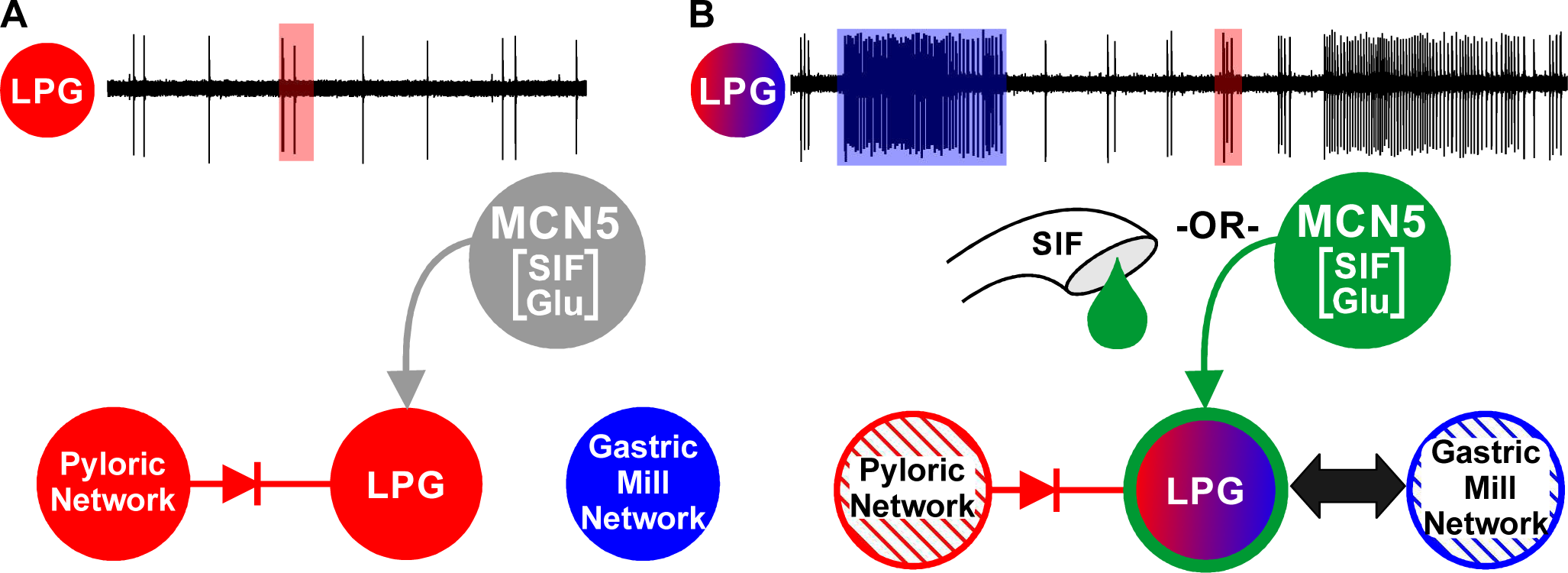
LPG switches from single-to dual-network activity via modulation of intrinsic properties. *A*, In control conditions, LPG participates in only the pyloric network (red), via rectifying electrical synapses with the pyloric pacemaker neurons. MCN5 (grey) is inactive *B*, Exogenously applied or endogenously released Gly^1^-SIFamide causes LPG to switch to dual-network participation via modulation of intrinsic properties (green outline) that enable gastric mill-timed bursting (red/blue), and electrical synapses that maintain pyloric-timed bursting (red). Red-hatched pyloric and blue-hatched gastric mill networks indicate that these are not necessary for LPG to burst in gastric mill time. Synaptic connections between LPG and gastric mill network neurons are likely modulated as well (black double arrow), but this has not yet been examined. Green arrow = modulatory input, diode = rectifying electrical coupling; SIF, Gly^1^-SIFamide; Glu, glutamate.

### Neuronal Switching

Network flexibility includes neuronal switching, in which neurons change their network participation via neuromodulatory inputs. This includes switching between networks, (Hooper and Moulins, 1990; Ramirez, 1998; Jean, 2001), merging multiple networks, or novel networks forming from participants of multiple networks (Dickinson et al., 1990; Weimann et al., 1991; Meyrand et al., 1994; Faumont et al., 2005).

Behaviorally-related networks are often co-active, and neurons can participate in multiple oscillatory networks simultaneously (Bucher et al. 2006; Jacobs et al. 2007; Larson et al. 1994; Rangel et al. 2016; Steriade et al. 1993; Weimann et al. 1991). However, dual-network activity is flexible, and thus single-to-dual-network switching also occurs (Dickinson, 1995; Gestreau et al., 2000; Tryba et al., 2008; Bartlett and Leiter, 2012; Schmidt and Martin Wild, 2014).

### Mechanisms of Neuronal Switching

Prior to this study, the identified mechanism for recruiting switching neurons into another network, or into dual-network activity, was modulation of synaptic strength. For example, in lobster STNS networks, different neuromodulatory inputs alter synaptic strength to recruit a pyloric neuron into the cardiac sac network (Hooper and Moulins, 1990), or to recruit pyloric, gastric mill, and oesophageal network neurons into one network to produce a novel rhythm (Meyrand et al., 1994).

Although synaptic modulation is essential to neuronal switching, modulation of intrinsic membrane properties can contribute to neurons leaving a network. For example, while synaptic modulation recruits STNS neurons into networks, modulation of intrinsic properties removes neurons from their home networks (Cazalets et al., 1990; Hooper and Moulins, 1990; Faumont et al., 2005). Additionally, intrinsic property modulation may play a primary role when an exogenous neuromodulator causes dual sigh/eupneic mouse respiratory neurons to leave the eupneic network, despite fast chemical synaptic transmission being blocked (Tryba et al., 2008). However, the inability to simultaneously monitor all, or completely isolate individual neurons in the relatively large respiratory network, raises the possibility that another mechanism, such as electrical synapse modulation, may underlie dual-to-single-network switching (Tryba et al., 2008). Further, intrinsic property modulation may serve a primary role in neuronal switching, even with unmodulated synapses (Drion et al., 2019).

In our study, it was possible that gastric mill network, or pyloric electrical synaptic input (albeit at a different frequency), or both, was necessary for switching LPG into dual-frequency oscillations. In a computational study, modulation of both electrical and chemical synaptic strength facilitates neuronal switching (Gutierrez et al., 2013). The small pyloric/gastric mill networks enabled us to determine that, for both exogenously-applied, and endogenously MCN5-released Gly^1^-SIFamide, LPG did not require synaptic input to generate voltage-dependent oscillations at a second frequency. Thus, intrinsic property modulation, without a synaptic component, enables a neuron to generate oscillations at a second frequency timed with a second network.

Modulating intrinsic instead of synaptic properties to enable dual-frequency bursting has potential consequences for network function. First, a strengthened synapse imposing another frequency onto a neuron implies a passive role of this switching neuron in the new network. However, when the switching neuron intrinsically generates the appropriate frequency bursting, it could contribute to rhythm generation in the new network. In control, LPG is part of the pyloric pacemaker ensemble (Marder and Bucher, 2007), whereas in the MCN5/Gly^1^-SIFamide state, it may contribute to rhythm generation for two networks, at two distinct frequencies. Second, when synaptic modulation “pulls” a switching neuron into another network, the enhanced synapse enables both appropriately-timed bursting and coordination with network neurons.

However, modulating intrinsic properties to elicit neuronal switching at the appropriate frequency is independent from coordination. In control, there are no functional synapses capable of coordinating LPG and gastric mill neurons (Marder and Bucher, 2007).

However, synapses in this and many other systems, are subject to extensive modulation (Harris-Warrick, 2011; Nadim and Bucher, 2014). Thus MCN5/Gly^1^- SIFamide likely modulate synapses along with LPG intrinsic properties. Separating the two components of neuronal switching enables independent regulation and may allow for switching neurons to actively contribute to coordination in the new network. We have not yet determined the modulatory actions responsible for coordination in the MCN5/Gly^1^-SIFamide gastric mill rhythm.

### Interactions between Synaptic and Intrinsic Properties

Although commonly known for synchronization, electrical synapses can have complex interactions with chemical synapses and intrinsic properties, and thus unexpected consequences for network output (Weaver et al., 2010; Connors, 2017; Jing et al., 2017; Marder et al., 2017; Nadim et al., 2017; Alcamí and Pereda, 2019). For example, electrical coupling enables rapid switches in network output among neurons connected via chemical synapses (Bem et al., 2005), a larger range of network outputs, and switching between networks oscillating at distinct frequencies (Gutierrez and Marder, 2014). Here, electrical coupling maintaining LPG activity in the pyloric network must be balanced with slow burst intrinsic currents eliciting periodic LPG escapes from the pyloric network. This balance may require MCN5/Gly^1^-SIFamide to decrease LPG-PD/AB coupling strength. Similar to chemical synapses, electrical synapses are highly modulated, including direct modulation of gap junction channels (Zsiros and Maccaferri, 2008; Lane et al., 2016), or indirect effects on functional coupling through modulation of ionic conductances (Szabo et al., 2010; Haas and Landisman, 2012; Nadim et al., 2017). Either MCN5 co-transmitter, Gly^1^-SIFamide or glutamate (Blitz et al. 2019; this study), could modulate electrical synapses. Identified MCN5-released glutamate actions involve fast ionotropic inhibition, however metabotropic glutamate receptors occur in STG neurons (Krenz et al., 2000), and modulate electrical synapses in other systems (Curti and O’Brien, 2016). Although biogenic amine modulation of electrical coupling is well-described (Johnson et al., 1994; Zsiros and Maccaferri, 2008; Szabo et al., 2010), peptide regulation of electrical transmission does occur (Wang et al., 2014; Cachope and Pereda, 2015). Such modulatory actions may directly alter LPG gap junctions or change the functional impact of LPG-PD/AB coupling due to MCN5/Gly^1^-SIFamide modulation of intrinsic properties underlying LPG slow bursting.

It is also possible that gap junction properties facilitate interactions between electrical synapses and intrinsic properties important for switching. For instance, pyloric pacemaker electrical synapses are rectifying, with depolarizing current flowing preferentially from PD/AB to LPG, and hyperpolarizing current from LPG to PD/AB neurons (Fig. 1B) (Shruti et al., 2014). Thus, intrinsic current(s) initiating LPG slow bursts may overcome electrical coupling because the rectification decreases electrical synaptic current as LPG depolarizes above PD/AB membrane potentials. Contributions of rectification to network output are likely underappreciated, as its identification requires directly recording from, and manipulating coupled neurons. However, they can play important functions within circuits, such as regulating network sensitivity to chemical synaptic input (Gutierrez and Marder, 2013; O’Brien, 2014; Palacios-Prado et al., 2014; Szczupak, 2016). Although rhythmic electrical synaptic input was not necessary for LPG slow intrinsic oscillations, with both networks intact, this input may regulate LPG slow bursting.

### Functional Implications of Neuronal Switching

Rhythmic behaviors such as respiration and locomotion, swallowing and vocalization, and feeding-related behaviors require coordination (Bernasconi and Kohl, 1993; Wood et al., 2004; Bartlett and Leiter, 2012). In fact, there are dangerous consequences to poor coordination, such as aspiration with improper swallowing/breathing coordination (Bartlett and Leiter, 2012). Furthermore, individual muscles can alternate between different rhythmic behaviors, or contribute to multiple simultaneous behaviors (Lancaster et al., 1995; Ramirez, 1998; Gestreau et al., 2000; Schmidt and Martin Wild, 2014). However, the extent of coordination varies based on behavioral version or environmental conditions (Lancaster et al., 1995; Stickford and Stickford, 2014; Stein and Harzsch, 2020). Neuronal switching between single- and dual-network activity may underlie such flexibility for network coordination.

Beyond rhythmic motor networks, systems-level studies suggest that neuromodulatory inputs elicit rapid reconfiguration of oscillatory networks involved in sensory and cognitive processes (Draguhn and Buzsáki, 2004; Roopun et al., 2008; Haider and McCormick, 2009; Ainsworth et al., 2011; Akam and Kullmann, 2014; Rangel et al., 2016). Specifically, a suggested major role of noradrenergic locus coeruleus (LC) neurons is large-scale, brain-wide network reconfiguration (Bouret and Sara, 2005; Grella et al., 2019), supported by fMRI studies in mice and humans (Hermans et al., 2011; Zerbi et al., 2019). In non-motor networks, it can be difficult to identify behavioral consequences of changing co-activation across brain regions.

Potential functions of rapid network reorganization, including switching between single and multiple oscillatory frequencies, include combining multimodal aspects of sensory perception, or multiple components of a memory for storage (Roopun et al., 2008; Wang, 2010; Akam and Kullmann, 2014). Dysfunction in these processes may lead to neurological disorders, suggested by decreased synchrony between brain regions in schizophrenia patients, and brain network dysregulation, potentially due to LC changes, in autism spectrum disorders (Uhlhaas and Singer, 2010; London, 2018). Given the many similarities in network function and plasticity from small to large networks (Haider and McCormick, 2009; Koch et al., 2011; Bargmann and Marder, 2013; Kopell et al., 2014), similar basic principles of neuronal switching in small networks will likely extend to larger, more diffuse networks.

## Acknowledgements

Funding from National Science Foundation IOS:1755283 (DMB). We thank Dirk Bucher for assistance with spectral analysis and Ryan Snyder for helpful discussions and feedback on the manuscript.

